# Treatment of prostate cancer with CD46 targeted ^225^Ac alpha particle radioimmunotherapy

**DOI:** 10.1101/2022.10.13.512165

**Authors:** Anil P. Bidkar, Sinan Wang, Kondapa Naidu Bobba, Emily Chan, Scott Bidlingmaier, Emily A. Egusa, Robin Peter, Umama Ali, Niranjan Meher, Anju Wadhwa, Suchi Dhrona, Denis Beckford-Vera, Yang Su, Ryan Tang, Li Zhang, Jiang He, David M. Wilson, Rahul Aggarwal, Henry F. VanBrocklin, Youngho Seo, Jonathan Chou, Bin Liu, Robert R. Flavell

## Abstract

Radiopharmaceutical therapy is changing the standard of care in prostate cancer (PCa) and other malignancies. We previously reported high CD46 expression in PCa and developed an antibody-drug conjugate and immunoPET agent based on the YS5 antibody, which targets a tumor-selective CD46 epitope. Here, we present the preparation, preclinical efficacy, and toxicity evaluation of [^225^Ac]DOTA-YS5, a radioimmunotherapy agent based on the YS5 antibody. Our radiolabeled antibody retains binding efficacy and shows a high tumor to background ratio in PCa xenografts. Furthermore, we show that radiolabeled antibody was able to suppress the growth of cell-derived and patient-derived xenografts, including PSMA-positive and deficient models. Nephrotoxicity, not seen at low radioactive doses, is evident at higher radioactivity dose levels, likely due to redistribution of daughter isotope ^213^Bi. Overall, this preclinical study confirms that [^225^Ac]DOTA-YS5 is a highly effective treatment and suggests feasibility for clinical translation of CD46 targeted radioligand therapy in PCa.

## Introduction

Prostate cancer (PCa) is the most diagnosed non-cutaneous malignancy and second leading cause of death in men worldwide in the year 2021.^1^ While early-stage tumors may be effectively treated with surgery, radiation, or androgen deprivation therapy, a significant fraction of patients progress to metastatic, castration-resistant prostate cancer (mCRPC), which is refractory to these treatments. Therapeutic options for patients with mCRPC include androgen signaling inhibitors, chemotherapy, immunotherapy, and radiopharmaceutical therapies including ^177^Lu-PSMA-617 (Lu-177 vipivotide tetraxetan) or ^223^Ra. Those treatments are not curative and patients eventually progress. As such, there is a need for preclinical and translational development of new treatments.

In PCa, prostate-specific membrane antigen (PSMA)-targeted radiopharmaceutical therapies have overwhelmingly attracted the most attention. PSMA is a type II glycoprotein overexpressed on PCa cells. Currently, PSMA PET (positron emission tomography) imaging using ^68^Ga-PSMA-11 or ^18^F-DCFPyL is used in routine clinical care to localize and stage patients. However, a significant fraction of mCRPC patients are PSMA-negative at diagnosis or develop PSMA negativity on imaging over the course of treatment, including in patients with treatment-emergent neuroendocrine/small cell PCa (t-SCNC).^2–5^ Immunohistochemistry analysis of primary and metastatic prostate cancer samples revealed that 7/51 primary tumors and 6/51 metastatic lesions had heterogeneous PSMA expression, and a further 2/51 primary tumors and 8/51 metastatic lesions were frankly negative.^6^ These prior studies demonstrate potential limitations of PSMA-directed imaging and radiopharmaceutical therapies, highlighting the need to identify and evaluate other targets in mCRPC.

We have previously used a non-gene expression based approach to identify novel tumor cell surface targets including conformational and post-translationally modified epitopes. We selected, by laser capture microdissection, billion-member phage human antibody display libraries on PCa patient specimens following counter selection on normal cells and tissues, and identified a panel of human antibodies binding to tumor selectively *in vitro* and *in vivo*.^7–9^ Following immunoprecipitation and mass spectrometry analysis, we identified one of the target antigens as CD46.^10^ CD46 is best known for being a negative regulator of the complement activation process.^11^ We showed that CD46 maintains a high level of expression in PCa across differentiation patterns.^10^ Compared with PSMA, CD46 is more homogeneously expressed especially in patients who become resistant to androgen blockade including t-SCNC.^10^ We developed and optimized a CD46 targeting human antibody, YS5, which binds to a tumor selective conformational epitope, enabling therapeutic targeting of CD46. We have developed an antibody-drug conjugates (ADC) based on YS5, demonstrated its preclinical efficacy in xenograft models, and shown that it is well tolerated in non-human primates.^10^ The ADC (FOR46) is being tested in mCRPC patients, as a single agent (phase I, NCT03575819) and a combination treatment (phase I/II, NCT05011188), with promising early results reported.^12^ To enable patient stratification, we have also developed an immunoPET agent based on YS5, ([^89^Zr]DFO-YS5), and shown that it detects adenocarcinoma, NEPC, and PSMA-positive or negative tumor xenografts in preclinical studies.^13^ A first-in-human study of this PET agent is ongoing in mCRPC patients (NCT05245006).

Radioimmunotherapy (RIT) exploits target-specific affinity agents to deliver therapeutic radioisotopes specifically to the target tissue. By using an antibody-based targeting approach, many RIT agents have been produced for clinical applications. Alpha-particle therapies demonstrate both potential advantages as well as limitations when compared to other radiopharmaceutical therapy methods, including greater efficacy, but also at the cost of potential increased toxicity. Actinium-225 (^225^Ac), an alpha-emitting radionuclide, is being explored for PCa and other malignancies.^14^ The advantage of using ^225^Ac is that it has a long half-life of 9.9 days, and it decays by emitting net four alpha particles generated from ^221^Fr (6.3 MeV), ^217^At (7.1 MeV), ^213^Bi (5.8 MeV), and ^213^Po (8.4 MeV). Early clinical studies in mCRPC show that ^225^Ac-PSMA-617 has efficacy, although treatment is associated with xerostomia, a target-dependent toxicity, and the daughter isotopes are believed to accumulate in kidneys, causing renal toxicity.^15^ A number of investigators are evaluating ^225^Ac labeled small molecules or antibodies for PCa in both preclinical and clinical studies.^16,17^ These studies reveal potential for ^225^Ac-based radiopharmaceutical therapy in PCa, although none of the agents have yet emerged as clinical standard of care. Herein, we report the development and preclinical evaluation of an alpha-particle therapy against CD46 by labeling the anti-CD46 antibody (YS5) with ^225^Ac ([^225^Ac]DOTA-YS5).

## Results

### CD46 is more persistently expressed in PCa patient-derived xenografts compared with PSMA, and may be imaged using immunoPET

Western blot analysis of the series of patient-derived xenografts (PDX) was carried out to study the expression of CD46 and PSMA. For this study, we utilized well-characterized patient-derived xenografts (PDXs) from the Living Tumor Laboratory.^18–20^ As shown in **Fig 1a** most of the xenografts, including all the neuroendocrine prostate models used in this experiment were PSMA-negative, but most showed high expression of CD46. Our previous study reported that PSMA-positive and PSMA-negative xenografts could be readily detected with a CD46 PET scan using [^89^Zr]DFO-YS5 probe, including the LTL-331, LTL-331R, and LTL-545 models.^13^ Herein we further demonstrate high uptake in the LTL-484 adenocarcinoma model, calculated at 23.7 ± 4.7% ID/g based on region of interest analysis of microPET/CT images (**Fig 1b**). Taken together with our prior report, these data demonstrate that CD46 is highly expressed across a panel of clinically relevant PDX, and a high amount of [^89^Zr]DFO-YS5 can be delivered and detected using PET imaging.

**Figure 1:**
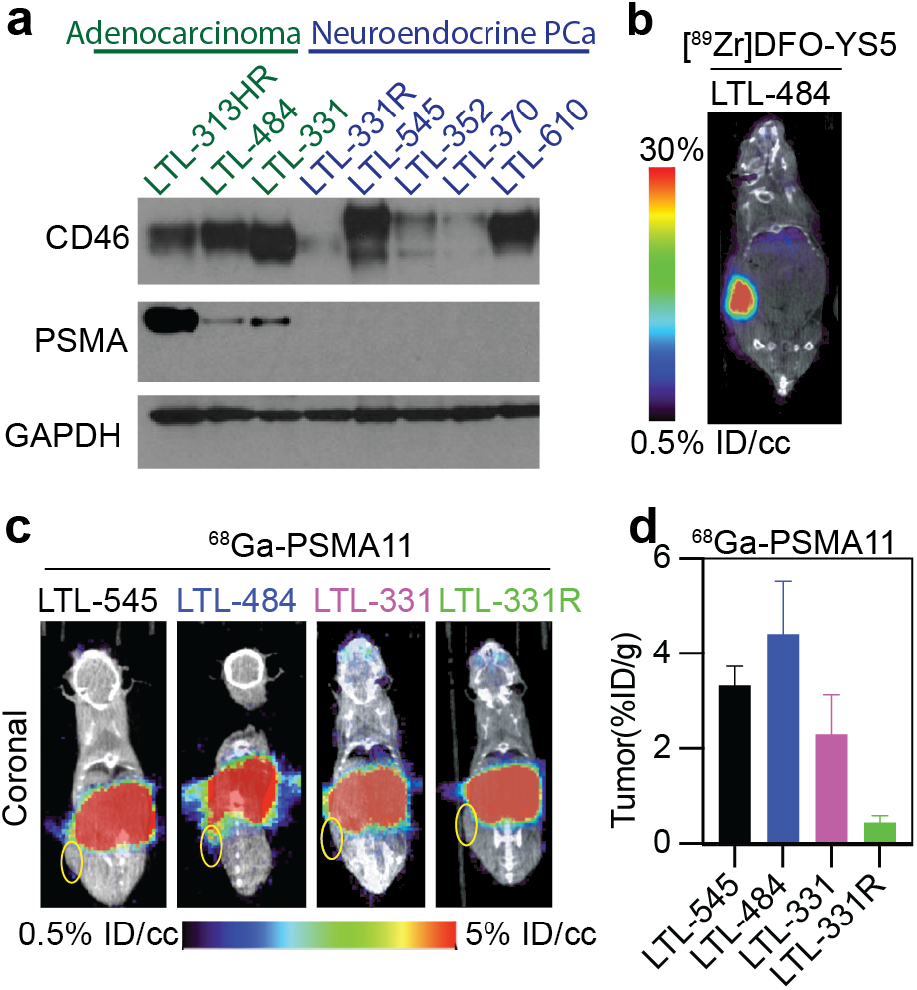
CD46 is a target for therapy in PCa, including in PSMA-negative disease. **a** Western blot analysis of the PDX tumors showing overexpression of CD46 in PSMA-positive as well as PSMA-negative tumors. **b** micro PET/CT image for detection of the LTL-484 model with the [^89^Zr]DFO-YS5 probe. **c** Coronal micro PET/CT images for LTL-545, LTL-484, LTL-331, and LTL-331R using ^68^Ga-PSMA-11. Tumor regions are shown by yellow ellipses. **d** Comparison of the tumor uptake of ^68^Ga-PSMA-11 in PDX model shows low to moderate uptake of ^68^Ga-PSMA-11 in the indicated PDX models.

To compare the results of CD46 targeting [^89^Zr]DFO-YS5 with a clinically used PSMA targeting probe, we performed micro PET/CT scans and biodistribution (**Fig 1c & d, Table S1**) using ^68^Ga-PSMA11 in LTL-545, LTL-484, LTL-331, and LTL-331R PDXs. Tumor uptake of ^68^Ga-PSMA11 was low to moderate; %ID/g for LTL-545, LTL-484, LTL-331, and LTL-331R was found to be 3.32±0.40, 4.41±1.11, 2.29±0.82, and 0.43±0.13, respectively (**Fig 1d**). The uptake seen in the LTL-545 and LTL-331R model may be attributable to non-PSMA mediated binding, given the low expression seen on western blotting analysis. These data suggest that CD46 is an actionable target in prostate cancer for imaging and therapy using the YS5 antibody, including in PSMA-low and negative tumors.

### Conjugation of DOTA to YS5 and ^225^Ac Radiolabeling

We adopted a standard approach for antibody radiolabeling and characterization utilizing DOTA chelator and ^225^Ac and characterized the conjugated antibody activity utilizing a recently developed immunoreactivity assay (**Fig 2a**).^21^ Firstly, the conjugation of DOTA to the YS5 antibody was performed in 0.1 M Na_2_CO_3_/NaHCO_3_ buffer, followed by size exclusion chromatography (SEC) purification. The successful conjugation and number of DOTA chelator on YS5 was confirmed with Matrix-assisted laser deposition/ionization-time of flight (MALDI-TOF) mass spectroscopy (**Fig S1a**), which revealed that an average of 8.7±0.2 (n=2) equivalents of DOTA were conjugated to each YS5. The DOTA-YS5 was stored at -20°C until radiolabeling reactions were carried out. The isolated yield of [^225^Ac]DOTA-YS5 from [^225^Ac]Ac(NO_3_) was 39.4±4.4% (n=6) with a specific activity of 0.10±0.01 µCi per µg of antibody (n=6). As shown in iTLC in **Fig 2b**, purity of radiolabeled antibody was 94.8±8.1% (n=3). In addition, size exclusion chromatography analysis (**Fig S1b**) showed no aggregation of the [^225^Ac]DOTA-YS5 after the labeling and purification process. *In vitro* stability of the radioimmunoconjugate, [^225^Ac]DOTA-YS5 was evaluated in human serum and saline buffers at 37°C for 14 days. As shown in **Fig S1c**, >85% of radioimmunoconjugate was intact at all the time points except for the 14^th^ day measurement in the saline buffer (∼55.24% is intact). The beads-based binding assay ^22^ results shown in **Fig 2c** confirm that binding of [^225^Ac]DOTA-YS5 was 100±4%, which was reduced to 25.6±1.5% in the blocking sample where 10-fold excess of cold (unlabeled) YS5 was added, and 14.9±10.6% in non-antigen coated beads. Overall, these results demonstrate that [^225^Ac]DOTA-YS5 was efficiently radiolabeled with retention of immunoreactivity.

**Figure 2:**
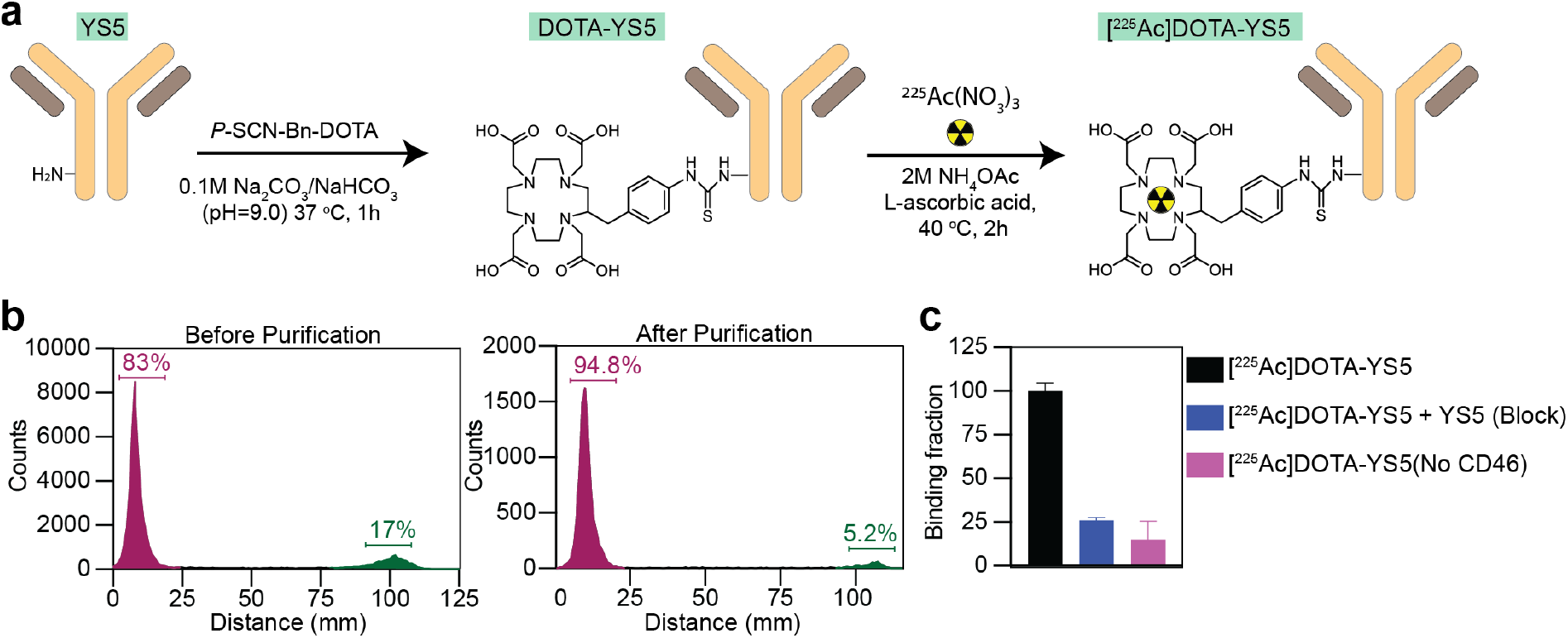
Preparation and confirmation of retention of immunogenicity of the [^225^Ac]DOTA-YS5 by *in vitro* studies. **a** Synthesis scheme of [^225^Ac]DOTA-YS5. **b** iTLC plots of the crude reaction and the purified [^225^Ac]DOTA-YS5 showing 94.84±8.1% (n=3) purity. **c** Magnetic beads assay showing 100±4% binding of [^225^Ac]DOTA-YS5 denoting the preservation of immunogenicity of radiolabeled antibody. Addition of cold YS5 reduces binding to 25.6±1.5%, demonstrating specificity.

### [^225^Ac]DOTA-YS5 demonstrates binding and cytotoxicity to PCa cells *in vitro*

Next, we studied binding and cytotoxicity of [^225^Ac]DOTA-YS5 to PCa cells *in vitro*. For binding, 22Rv1 cells incubated with [^225^Ac]DOTA-YS5 at 7 nM showed 19.25±0.70% binding (**Fig S2**), which was reduced to 4.46±0.17% by the addition of 2-fold cold YS5. The *in vitro* uptake of [^225^Ac]DOTA-YS5 was studied on 22Rv1 and DU145 cells. Results in **Fig 3a** and **3b** show number of cell-associated (membrane-bound, internalized, and total) molecules for the cells incubated with [^225^Ac]DOTA-YS5. Interestingly, the two cell lines used in this study show different binding and internalization patterns. In case of 22Rv1 cells (**Fig 3a**), most of the [^225^Ac]DOTA-YS5 was internalized or bound in the first 30 minutes of incubation; whereas the membrane-bound activity increased up to 24 h. Total cell-associated molecules (membrane-bound + internalized) for 22Rv1 cells at 24 h were 9.7×10^11^±4.5×10^10^, which suggests that a greater number of [^225^Ac]DOTA-YS5 molecules were interacting with CD46 on cell surface over time. In contrast, the total number of [^225^Ac]DOTA-YS5 antibody molecules bound to DU145 cells (**Fig 3b**) increased from 8.0×10^10^±7.6×10^9^ at 30 minutes to 4.5×10^11^±1.9×10^9^ molecules at 24 h.

**Figure 3:**
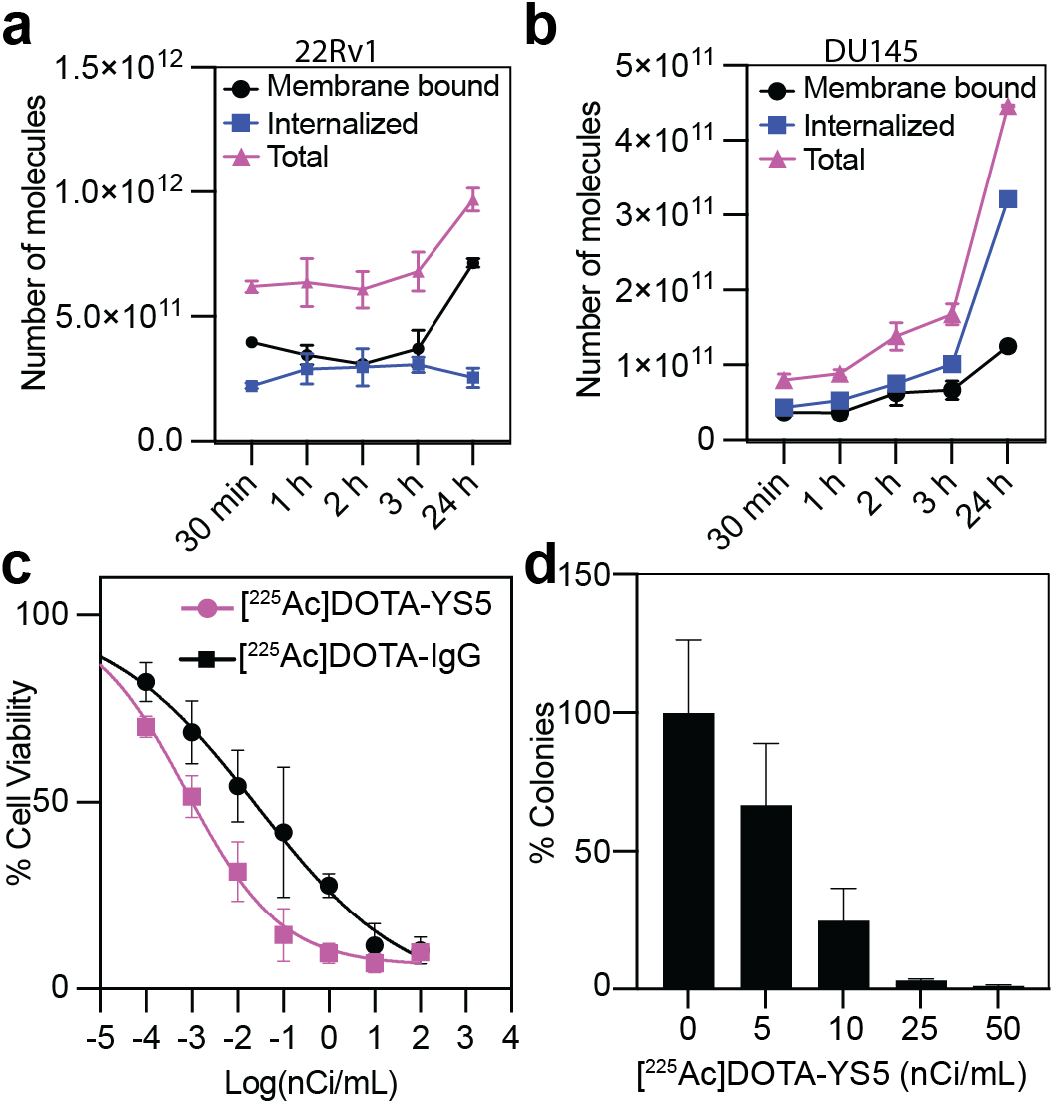
[^225^Ac]DOTA-YS5 shows high binding efficiency, internalization, and cell killing in *in vitro* assays. **a, b** Cell associated molecules of [^225^Ac]DOTA-YS5 in 22Rv1 (**a**) and DU145 (**b**) cells after the treatment for different time points. **c** MTT assay showing dose-dependent reduction of cell viability from [^225^Ac]DOTA-YS5 treatment on 22Rv1 cells (IC_50_: 80.0 ±20.0 pCi/mL for [^225^Ac]DOTA-YS5 and 7.3 ±0.9 nCi/mL for [^225^Ac]DOTA-IgG). **d** Clonogenic survival of the 22Rv1 cells treated with [^225^Ac]DOTA-YS5 showed a dose-dependent decrease in the number of colonies after treatment.

The cell-killing activity of [^225^Ac]DOTA-YS5 was studied with cell viability and clonogenic assays. A dose-dependent reduction in cell viability was observed by the treatment of [^225^Ac]DOTA-YS5 on 22Rv1 cells (**Fig 3c**). The half-maximal inhibitory concentrations (IC_50_) of [^225^Ac]DOTA-YS5 and control nonspecific [^225^Ac]DOTA-IgG were found to be 80.1±23.5 pCi/mL, and 7.34 ±0.9 nCi/mL (**Fig 3c**), respectively. In the clonogenic assay where tumor cells grow in 3D as opposed to monolayer, IC_50_ of [^225^Ac]DOTA-YS5 was determined to be 10.09±3.61 nCi/mL (**Fig 3d**). These data suggest that [^225^Ac]DOTA-YS5 has high specific binding and potent cytotoxicity against PCa cells.

### *In vivo* biodistribution, tumor uptake, and autoradiographic analysis of [^225^Ac]DOTA-YS5

Since imaging of [^225^Ac]DOTA-YS5 is challenging owing to low emission rate of imageable photons, we utilized *ex vivo* biodistribution studies to assay the tissue distribution of the radiopharmaceutical. Mice bearing 22Rv1 xenografts were administered 0.5 µCi doses of [^225^Ac]DOTA-YS5 via tail vein. As shown in **Fig 4a** and **Table S2**, tumor uptake of [^225^Ac]DOTA-YS5 at 24 h (Day 1) post injections (p.i.) was 11.64±1.37 %ID/g, which kept increasing up to 96 h (Day 4) to 28.58±10.88 %ID/g. The average %ID/g for 168 h (Day 7), 264 h (Day 11), and 336 h (Day 14) were found to be 29.35±7.76, 18.10±8.12, and 21.12±12.30, respectively (**Fig 4a**). Similarly, the last observation on 408 h p.i (Day 17) showed uptake of 31.78±5.89 %ID/g. The %ID/g for tumors was higher than non-targeted organs for all time points, except 24 h, where a high uptake was seen in kidneys. Tumor to blood, tumor to kidney, and tumor to muscle ratio (**Fig 4b, Table S3**) increased from day 1 to day 17, suggesting the clearance of the [^225^Ac]DOTA-YS5 in most of the organs including muscle and kidneys, with simultaneous accumulation in tumor tissue. As seen from gamma energy spectra in **Fig 4c** & **Fig S3**, the higher uptake seen in kidneys at 24 h could be due to the accumulation of ^213^Bi (from ^225^Ac decay) in the kidneys, a phenomenon that has been reported by others.^23,24^ This was not observed in other tissues, including the bones (**Fig S3**). The reduction in activity of blood from 24 h (11.64±1.37 %ID/g) to 408 h (1.71±0.47 %ID/g) indicates gradual clearance via liver and kidney (**Fig 4a**).

Digital autoradiographic imaging (iQID camera) was performed to study tissue distribution of [^225^Ac]DOTA-YS5 in tumor and mouse organs. **Fig 4d** shows iQID camera digital autoradiographs for organs collected on day 1, day 2, day 4, and day 7 after injections of [^225^Ac]DOTA-YS5. Signal from [^225^Ac]DOTA-YS5 was not uniform in tumor tissues (**Fig 4d**), possibly due to the presence of necrotic core cells, uneven vascular distribution, or heterogenous expression of CD46. Increased accumulation of radioactivity was observed in the renal cortex. For the remaining organs, distribution of [^225^Ac]DOTA-YS5 was homogenous.

**Figure 4:**
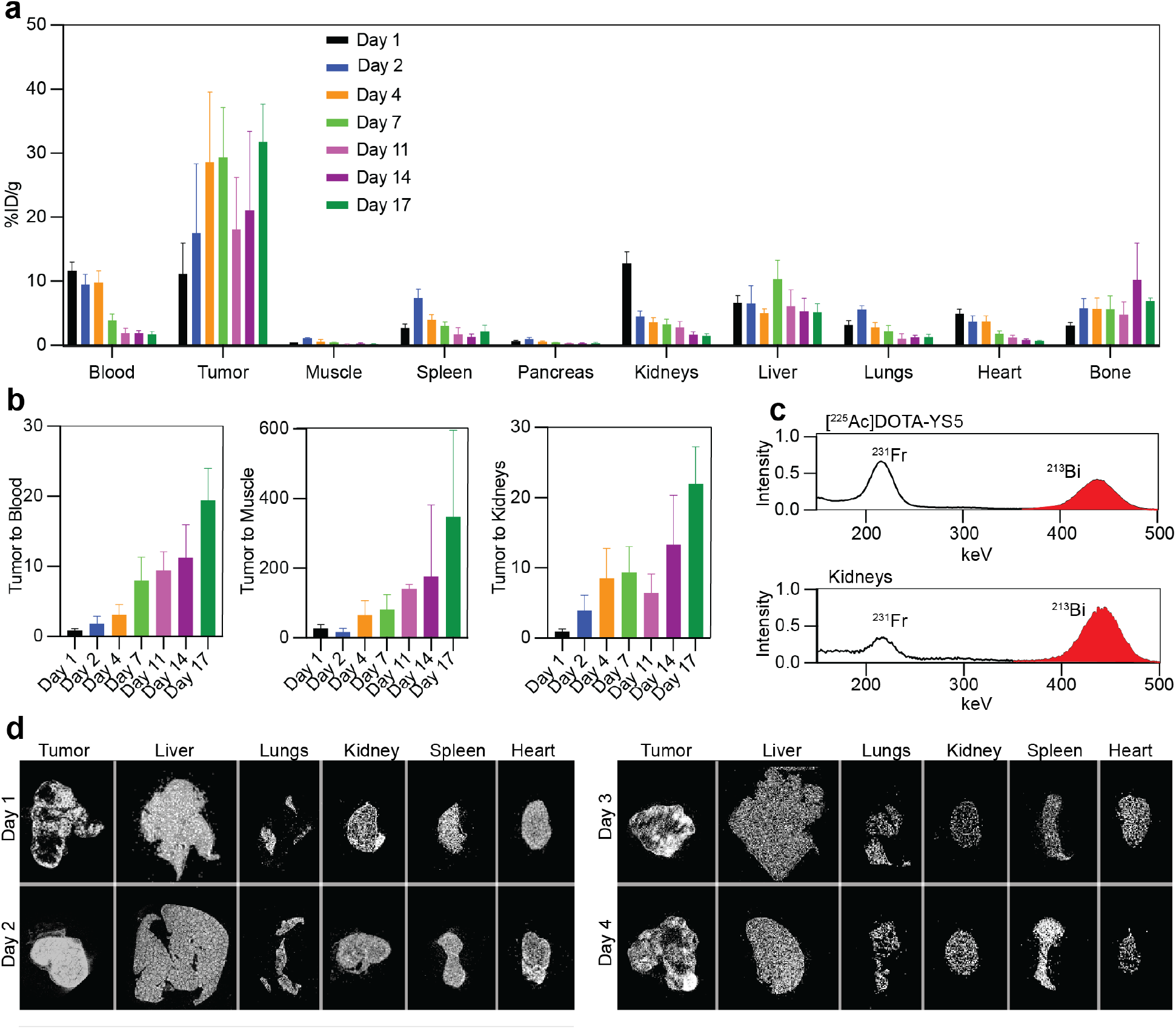
Biodistribution of the [^225^Ac]DOTA-YS5 in 22Rv1 xenograft bearing mice confirming high delivery of the [^225^Ac]DOTA-YS5 to tumor tissue. **a** The distribution of [^225^Ac]DOTA-YS5 (represented as %ID/g, mean ±SD, n=4) in organs collected from day 1 to day 17. **b** Tumor to blood, tumor to muscle, and tumor to kidney ratios from the biodistribution studies indicate [^225^Ac]DOTA-YS5 clearance with simultaneous accumulation in tumor tissue. **c** Gamma energy spectra showing accumulation of ^213^Bi in kidneys. As compared to the equilibrium gamma energy spectra of [^225^Ac]DOTA-YS5, increased intensity of the ^213^Bi was observed in kidney samples at 24 h. **d** iQID camera digital autoradiographic imaging for distribution of the [^225^Ac]DOTA-YS5 in 22Rv1 xenograft tumor and healthy tissues studied from day 1 to day 7 post injections. The distribution of the radioactivity in tumor tissue is heterogenous, whereas all the other organs show homogenous distribution.

### [^225^Ac]DOTA-YS5 treatment induces DNA double-strand breaks in tumor tissue

As shown in **Fig 5**, the 22Rv1 xenografts treated with 0.5 µCi dose of [^225^Ac]DOTA-YS5 were stained with hematoxylin and eosin (H&E) and a DNA damage marker protein (phosphorylated γ-H2AX). **Fig 5a, b** shows that treatment of 0. 5 µCi dose of the [^225^Ac]DOTA-YS5 for day 7 or day 14 increases the γ-H2AX foci in the cell nucleus confirming DNA double-strand breaks. The number of γ-H2AX foci per cell in 7 days treatment of 0.5 µCi doses were found to be 4.46±0.75, significantly higher than saline (1.37±0.10 foci per cell, p-value, 0.02 **Fig 5a, Fig S4**). Similarly, as shown in **Fig 5b** and **Fig S4**, a higher number of the γ-H2AX foci (4.90±0.55 per cell) were also observed for 14 days p.i. of [^225^Ac]DOTA-YS5 (p-value, 0.02). Additionally, DAPI and H&E staining for the tumors from [^225^Ac]DOTA-YS5 14 days post treatment (**Fig 5b**), showed deformed and pycnotic nuclei confirming high nuclear damage.

**Figure 5:**
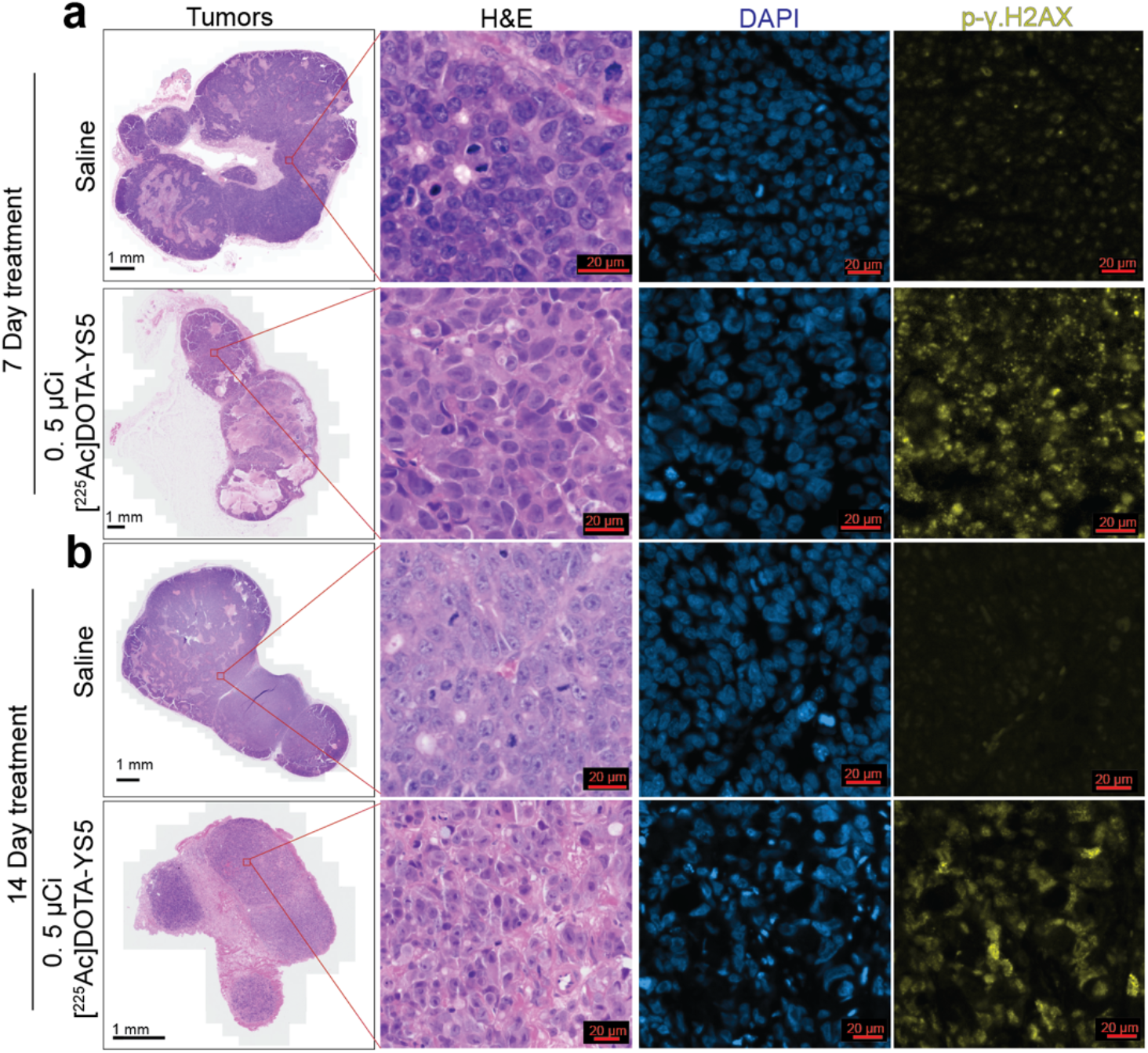
Histology and immunofluorescence imaging showing DNA damage after [^225^Ac]DOTA-YS5 treatment. **a**,**b** H&E and phosphorylated γ-H2AX (p-γ.H2AX) staining confirms morphological changes and p-γ.H2AX foci for [^225^Ac]DOTA-YS5 treatment for 7 days (**a**) and 14 days (**b**).

### Toxicity and Maximum tolerated dose

Initial acute toxicity studies were carried out for 15 days in male nude mice (without tumor) injected with saline, 0.06 µCi, 0.125 µCi, 0.25 µCi, and 0.5 µCi doses of [^225^Ac]DOTA-YS5. Bodyweight measurements shown in **Fig S5a** indicate no significant weight loss in [^225^Ac]DOTA-YS5 treated mice for 15 days. Following the 15 days of observation, mice in these groups were euthanized for further evaluation. The liver and kidney function test results shown in **Fig S5b, and Table S4** demonstrate no significant difference in saline vs. treated groups. Along with the liver and kidney function test, complete blood count measurements also showed no significant changes in the blood cell counts for saline vs. treated groups (**Fig S5c, Table S5**). Histologic evaluation of the organs collected after 15 days showed no significant changes in the saline vs. treated groups.

Following the acute toxicity studies, chronic toxicity studies were performed (**Fig 6**, Supplemental **Table S6, S7**). The male nude mice were injected with saline, 0.25 µCi, 0.5 µCi, or 1 µCi doses of [^225^Ac]DOTA-YS5 and were observed for 117 days for any sign of illness, pain, and body weight loss. All the mice in 1 µCi dose treatment group were euthanized on 14^th^ day p.i. due to significant toxicity and body weight loss (**Fig 6a**). Out of four animals in the 0.5 µCi dose group, a significant body weight loss was observed, and one mouse was euthanized on day 103 due to body weight loss by 20%, while the saline and 0.25 µCi dose demonstrated no detectable body weight loss (**Fig 6a, b**). On laboratory analysis, creatine and blood urea nitrogen levels were elevated at 0.5 µCi dose (**Fig 6c, Table S6**), consistent with renal toxicity. Alkaline phosphatase levels (for liver or bone damage) were also higher in 0.5 µCi dose treatment (**Fig 6c)**, while the liver function tests for alanine transaminase and aspartate transaminase were unchanged. There were no significant changes in blood counts for saline vs. treated groups (**Fig 6d, Table S7**).

**Figure 6:**
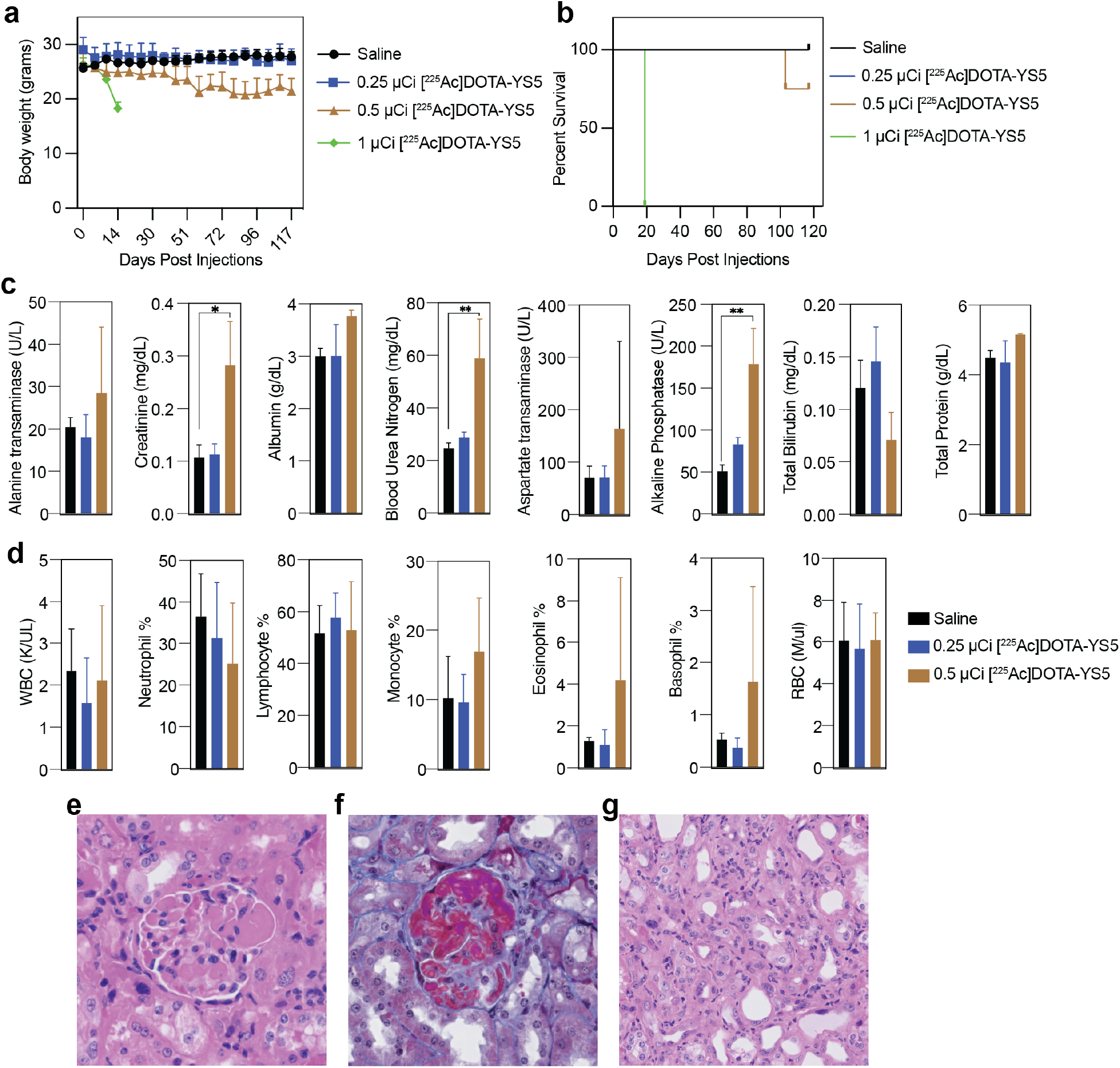
Chronic toxicity study of the [^225^Ac]DOTA-YS5 in nude mice. **a** The body weight measurements of the mice injected with the [^225^Ac]DOTA-YS5 show a gradual decrease with 0.5 µCi doses, whereas no significant body weight loss was seen in the 0.25 µCi dose or in the saline control group. **b** Survival plot in the toxicity study. **c** Liver and kidney function tests results showing increase in creatinine, blood urea nitrogen, and alkaline phosphatase. **d** Blood cell counts confirming no significant difference in saline control vs [^225^Ac]DOTA-YS5 injected mice. **e** Histologic H&E findings in kidney of a mouse injected with 0.5 µCi dose with diffuse parenchymal damage and occlusion of glomerular capillary loops by fibrin thrombi with glomerulosclerosis. **f** Trichrome stain highlighting fibrin thrombi. **g** Higher power images showing tubular injury in the mouse treated with [^225^Ac]DOTA-YS5. One-way ANOVA p values are indicated as * p < 0.05. ** p < 0.01, *** p < 0.001.

Histologic evaluation of the organs collected in this chronic toxicity study revealed that 0.5 µCi dose resulted in significant renal damage. As shown in **Fig 6e-g**, for 0.5 µCi dose, glomeruli demonstrated marked fibrin deposition in the capillary loops and glomerulosclerosis (**Fig 6e, f**). Additionally, the renal tubules showed dilatation and flattening with scattered cells containing enlarged nuclei with anisonucleosis and necrotic luminal epithelial cells **(Fig 6g)**. Other organs including spleen, bone and bone marrow, lungs, heart, and liver showed no histologic evidence of damage (**Fig S6**). Based on these studies, we conclude that the maximum tolerated dose was 0.25 µCi.

### [^225^Ac]DOTA-YS5 is effective in reducing tumor volume and prolonging survival in cell line-derived PCa models

CD46 expressing cell line-derived PCa xenograft models were used to study therapeutic efficacy of [^225^Ac]DOTA-YS5. Cohorts of n=7 22Rv1 xenograft (PSMA-positive) bearing mice were randomized to injection with saline, 0.25 µCi, 0.5 µCi doses of [^225^Ac]DOTA-YS5, or 0.5 µCi of [^225^Ac]DOTA-IgG. Non-targeting [^225^Ac]DOTA-IgG was kept as a negative control test to assay for nonspecific treatment effects. As shown in **Fig 7a**, treatment with 0.25 µCi and 0.5 µCi doses of [^225^Ac]DOTA-YS5 significantly inhibited tumor growth in a dose-dependent manner, while mice in control cohorts (saline and [^225^Ac]DOTA-IgG) showed rapid tumor growth. The survival plot (**Fig 7a**) indicates that median survival of the mice treated with saline and [^225^Ac]DOTA-IgG were 37 days and 51 days (p= 0.0012), respectively. In contrast, the median survival of 0.25 µCi and 0.5 µCi doses of [^225^Ac]DOTA-YS5 was significantly improved to 103 (p=0.0006) and 131 days (p=0.0003), respectively. The body weight measurements of the mice in these groups are shown in **Fig 7a**; results indicate that a 0.5 µCi dose of [^225^Ac]DOTA-YS5 results in a gradual decrease in body weights over the period, consistent with the toxicity studies. **Fig 7a** shows that 0.5 µCi dose of [^225^Ac]DOTA-YS5 has the highest antitumor response but also greater toxicity. Overall, these data demonstrate that 0.25 and 0.5 µCi doses of [^225^Ac]DOTA-YS5 are effective for treatment of 22Rv1 xenografts, with toxicity observed at the 0.5 µCi dose.

**Figure 7:**
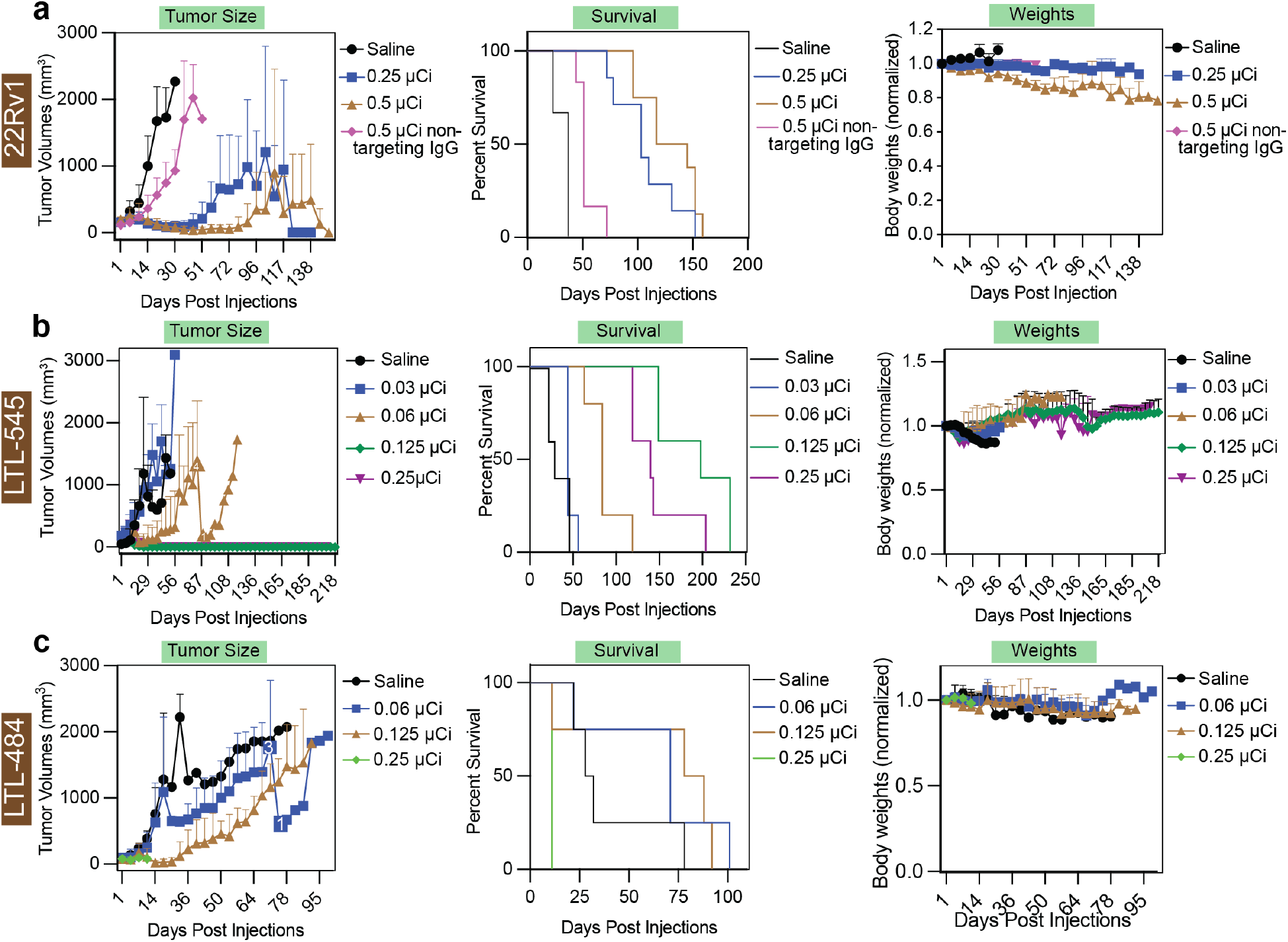
Antitumor activity of [^225^Ac]DOTA-YS5 in subcutaneous 22Rv1 and PDX tumor models. **a** Effect of [^225^Ac]DOTA-YS5 in 22Rv1 tumor-bearing mice. Tumor volume measurements demonstrated delayed 22Rv1 tumor growth in 0.25 µCi, and 0.5 µCi dose groups of [^225^Ac]DOTA-YS5. Kaplan-Meier survival plot showing improved survival probability of the [^225^Ac]DOTA-YS5 treated 22Rv1 xenograft bearing mice compared against control groups. Average body weights of the mice indicated gradual body weight loss in the 0.5 µCi treatment group. **b** Effect of [^225^Ac]DOTA-YS5 in LTL-545 PDX. Tumor volumes, overall survival, and average body weights in the animals administered with [^225^Ac]DOTA-YS5 at 0.03, 0.06, 0.125, and 0.25 µCi dose treatment. **c** Tumor size, overall survival, and body weight for the LTL-484 PDX mice treated [^225^Ac]DOTA-YS5.

We also tested a fractionated therapy regimen to further evaluate the treatment efficacy of [^225^Ac]DOTA-YS5. We administered fractionated dose (three doses) of 0.125 µCi of [^225^Ac]DOTA-YS5 to the mice with 22Rv1 xenografts on day 1, day 37, and day 49. As shown in **Fig S7**, the three fractionated doses of 0.125 µCi dose of [^225^Ac]DOTA-YS5 delayed tumor growth significantly compared to the saline control group. The median survival of the saline (17 days) was significantly lower than the study group treated with three doses of 0.125 µCi (71.5 days, p=0.0002)) of [^225^Ac]DOTA-YS5 (**Fig S7)**. Out of the ten mice in the fractionated dose group (0.125 µCi), no mouse was euthanized due to bodyweight loss, while only three mice showed a transient 15% body weight loss (**Fig S7**), and all mice regained their body weights. These data suggest that a fractionated therapy approach represents a potential method for treatment utilizing [^225^Ac]DOTA-YS5.

Additionally, a CD46 positive, PSMA-negative PCa cell line-derived xenograft model (DU145) was also employed to study the effect of [^225^Ac]DOTA-YS5. As shown in **Fig S8**, 0.25 µCi and 0.5 µCi dose of [225Ac]DOTA-YS5 had an antitumor effect that delayed the tumor growth. In fact, the 0.5 µCi dose showed tumor growth inhibition as soon as day 5 of the injections, and all mice from this group showed complete response. However, due to the indolent growth of the tumors in the control group in this study, an overall survival benefit was not observed (**Figure S8**).

### Antitumor effect of [^225^Ac]DOTA-YS5 in patient-derived xenograft models

We next evaluated antitumor potential of the [^225^Ac]DOTA-YS5 in PDX models. CD46 overexpressing patient-derived xenografts were used for studying the therapeutic efficiency of the [^225^Ac]DOTA-YS5. The PSMA-negative neuroendocrine prostate cancer PDX, LTL-545 bearing mice were injected with saline, 0.03 µCi, 0.06 µCi, 0.125 µCi, and 0.25 µCi dose of the [^225^Ac]DOTA-YS5 (**Fig 7b**). As shown in tumor volume plot (**Fig 7b)**, all the mice from 0.125 µCi, and 0.25 µCi groups demonstrated complete resolution of the tumor, whereas 0.06 µCi delayed the tumor growth as compared to saline control. Median survival of the animals treated with saline, 0.03 µCi, 0.06 µCi, 0.125 µCi, and 0.25 µCi were found to be 29 days, 44 days (not statistically significant), 84 days (p= 0.0025), 198 days (p= 0.0025), and 140 days (p= 0.0025), respectively (**Fig 7b**). During the first 15 days of treatment (**Fig 7b**), the body weights of LTL-545 bearing mice treated with [^225^Ac]DOTA-YS5 decreased, but these mice gradually regained their body weights.

NSG mice bearing LTL-484 PDX (CD46 and PSMA positive) were randomized into four arms for saline and [^225^Ac]DOTA-YS5 injections (0.06 µCi, 0.125 µCi, and 0.25 µCi) (**Fig 7c**). The 0.25 µCi dose injected mice demonstrated acute toxicity manifested by hunched posture and reduced locomotion on 11^th^ day post injection. On the other hand, we found that that 0.06 (median survival= 71, p= 0.335) and 0.125 µCi (median survival= 83, p=0.169) dose delayed the tumor growth with a non-statistically significant trend towards increase in the median survival as compared to saline control (median survival= 30). The NSG mice used in these experiments were more sensitive to radiation dose compared to nude mice used for the cell line derived xenograft studies, with a constant decrease in body weight seen over the period of treatment (**Fig 7c**). ^25^

## Discussion

Alpha-particle therapies hold great potential for treating metastatic prostate cancer. The PSMA-targeted alpha-particle therapy (^225^Ac-PSMA-617) showed potential antitumor response but with considerable toxicities.^26^ Recently, several studies have acknowledged the issue of less or no expression of PSMA in advanced prostate tumors which could result in resistance to PSMA directed therapies.^27^ Our prior report taken together with new data (**Fig 1**) showed that anti-CD46 antibody (YS5) based radiotracer probe ([^89^Zr]DFO-YS5) successfully targeted the overexpression of CD46 on PCa cell surface for PET imaging.^13^ In contrast to this, PSMA PET, as well as biodistribution studies with ^68^Ga-PSMA-11, show lower uptake in PDX tumors. Inspired by these results as well as prior studies demonstrating efficacy the YS5 antibody as an antibody-drug conjugate, we hypothesized that the YS5 antibody would serve as an ideal targeting vector for delivery of therapeutic alpha particles.^10^

We report the CD46 targeted alpha-particle therapy with [^225^Ac]DOTA-YS5 for PSMA-positive and PSMA-negative tumors. We developed a reproducible radiosynthesis and found that [^225^Ac]DOTA-YS5 was able to kill the PCa cell at higher efficiency as compared to the non-targeted [^225^Ac]DOTA-IgG control.^28^ In a biodistribution study high tumor uptake of [^225^Ac]DOTA-YS5 was observed over 17 days, along with clearance from healthy organs. We noted all the healthy organs except liver and bones demonstrated gradual clearance of [^225^Ac]DOTA-YS5. The primary route for metabolism of large molecules is via the liver; this could be the reason that no gradual decrease in activity was seen from liver.^29^ The steady uptake in liver corroborates with previous studies reported for ^225^Ac labeled PSMA-targeting antibodies.^30^ Similar results for liver were also noted in earlier studies with [^89^Zr]DFO-YS5 over the period of 168 h.^13^ Favorable tumor to blood and tumor to muscle ratios were found, suggesting feasibility for therapeutic studies.

We conducted a detailed toxicologic analysis following treatment, with the key finding of dose limiting nephrotoxicity. Our data suggest that this is due at least in part to non-target mediated redistribution of the daughter isotope, ^213^Bi. Though an antibody-chelate may retain the ^225^Ac, as the time passes ^225^Ac decays into daughter isotopes ^213^Bi and ^221^Fr which are ejected from the chelator due to the recoil effect. These daughter isotopes are reported to accumulate in kidneys, resulting in off-target toxicity.^24^ Our observations from the gamma-ray spectra of ^213^Bi and ^221^Fr confirm that there is a non-homogenous distribution of daughter nuclei in mouse organs. Specifically, comparison of gamma spectra between organs in **Fig S3** shows increased ^213^Bi peak intensity and therefore ^213^Bi accumulation in kidneys. Additionally, histopathology analysis reveals that ^213^Bi accumulation results in renal parenchymal damage, fibrin thrombi formation, and glomerulosclerosis. We conducted a dosimetry estimation, neglecting the daughter ^213^Bi, which did not demonstrate disproportionate renal radiation dose, further supporting our hypothesis that redistribution plays a key role (**Fig S9**). This substantiates the previous finding of ^213^Bi accumulation in kidneys resulting in kidney toxicity. Unexpectedly, we have also observed uptake of [^225^Ac]DOTA-YS5 in bone, but no damage to bone marrow or hematological toxicity was seen, possibly due to the short-range and low penetrability of alpha particles.

Evaluation of therapeutic efficiency revealed that [^225^Ac]DOTA-YS5 shows a significant antitumor response across a panel of PCa models, including in 22Rv1 (weakly PSMA-positive) and DU145 (PSMA-negative), as well as the LTL-545 (PSMA negative neuroendocrine) and LTL-484 (PSMA positive adenocarcinoma) patient-derived xenograft models. Marked increases in overall survival were observed in the 22Rv1 and LTL-545 models, with the latter demonstrating complete eradication of the tumor at the higher dose levels with no evidence of recovery even at delayed time points. Considering the long period of therapeutic efficiency studies, 0.25 µCi was the maximum tolerated dose (MTD) in the nu/nu model, with a prominent antitumor response. In the highly radiosensitive NSG model (required to grow the PDX models), the MTD is at a lower dose (0.125 µCi) with significant anti-tumor effects and an increase of survival.

Apart from PSMA, potential targets for PCa include prostate stem cell antigen (PSCA) and Delta-like ligand 3 (DLL3), kallikrein 2 (hK2), and CUB Domain-Containing Protein 1 (CDCP1). In preclinical studies, alpha particle emitter (^211^At) and beta emitter (^131^I and ^177^Lu) labeled PSCA targeted antibody fragments have shown tumor growth inhibition which significantly extended median survival in treated groups.^31,32^ Similarly, Human kallikrein peptidase 2 (hK2) targeted ^225^Ac labeled antibody showed excellent tumor targeting ability (VCaP 52.2 %ID/g and LNCaP-AR 24.5 %ID/g) and DNA damaging effect. Additionally, DNA damage events further increase the expression of hK2, resulting in feed-forward alpha particle therapy.^16^ Recently, Zhao *et al*. confirmed the expression of CUB Domain-Containing Protein 1 (CDCP1) on PSMA-negative metastatic CRPC tumors, followed by radiotracer probe and radioligand therapy development for targeting CDCP1.^33^ Due to the loss of PSMA and SSTR on NEPC, these tumors cannot be treated with PSMA or SSTR targeted RLTs. Such NEPC can be detected or treated by targeting novel cell surface marker as delta-like ligand 3 (DLL3).^34^ Findings from these preclinical and clinical studies with different radioisotopes and targeting molecules show promising results. Our results demonstrate that CD46 targeting with [^225^Ac]DOTA-YS5 could deliver the radiopharmaceutical to a broader range of prostate cancers compared with PSMA targeting.

Feasibility of translation of [^225^Ac]DOTA-YS5 to the clinic is high. We discovered that CD46 is an excellent target for mCRPC across differentiation patterns, developed a human antibody YS5 that binds to a tumor selective conformational epitope, and constructed an YS5-based ADC, ^10^ which is in multiple clinical trials (NCT03575819 and NCT05011188). In addition, we recently developed an YS5-based PET agent,^13^ which is now in a first-in-human study in mCRPC patients (NCT05245006). Thus, there is an established path for YS5-based agents to move from the bench to the clinic.

Safety is a critical issue that needs to be carefully investigated. While ^225^Ac-antibodies have demonstrated great efficacy, prior studies have also revealed dose-dependent toxicity.^17,35^ Potential mechanisms for toxicity include redistribution of the daughter isotopes away from the original site, due to the “recoil” effect.^36,37^ We observed this effect in higher uptake of ^213^Bi in the kidney compared to the other organs. Our results are consistent with prior literature which demonstrates both remarkable efficacy but also toxicity for treating cancer with ^225^Ac radiopharmaceutical therapy. There are several potential routes to mitigate this toxicity, including dose reduction, chelation therapy, linker optimization, or pretargeting.^38–40^

In conclusion, we have prepared and developed [^225^Ac]DOTA-YS5 as a radioimmunotherapy agent for the treatment of PCa, which shows efficacy for both PMSA deficient and PSMA-positive tumors. Due to the CD46-targeting nature of [^225^Ac]DOTA-YS5, we were able to deliver a high amount of radiation dose to tumor tissue, resulting in a potent antitumor effect. The data presented here provide strong evidence of therapeutic efficacy of [^225^Ac]DOTA-YS5 in preclinical models, and support future clinical translation of CD46-targeted radioligand therapy.

## Methods

### Western blotting

The PDX tissues were lysed in radioimmunoprecipitation assay (RIPA) buffer with protease and phosphatase inhibitor cocktail (Sigma-Aldrich: #P8340). The protein concentrations were calculated by BCA assay, and proteins from all the lysates were separated on SDS-PAGE (sodium dodecyl sulfate-polyacrylamide gel electrophoresis). Finally, these proteins from the gel were transferred to PVDF membrane, blocked (with 5% BSA), stained with primary antibodies (Cell Signaling Technology: anti-CD46, #13241; anti-PSMA #12815; and anti-GAPDH #2118), and were detected with HRP-conjugated secondary antibody along with the chemiluminescent substrate.

### ^68^Ga-PSMA11 imaging and biodistribution

^68^Ga-PSMA11 PET scans and biodistribution studies were performed to study the ability of ^68^Ga-PSMA11 to detect the patient-derived xenograft tumors. ^68^Ga-PSMA11 was synthesized according to the previously reported method using ^68^Ga/^68^Ga generator.^41^ For this, LTL-545, LTL-484, LTL-331, and LTL-331R PDX bearing mice were injected 100-160 μCi with ^68^Ga-PSMA11, and the PET scans were performed 1-4 h post injections similar to our prior reported methods.^42^ Following the PET scans, tumors and selected organs were collected, and the activity of ^68^Ga-PSMA11 was counted in the HIDEX gamma counter.

### [^89^Zr]DFO-YS5 imaging on LTL-484 PET and biodistribution

The labeling of ^89^Zr on DFO conjugated YS5 was done according to our previously reported protocol.^13^ The LTL-484 tumors-bearing mice were intravenously co-injected with 100-110 μCi (100μL volume) of [^89^Zr]DFO-YS5 and cold IgG (0.5 mg per mouse, Abcam ab91102).

### Antibody conjugation and ^225^Ac radiolabeling

The YS5 human IgG1 antibody was produced in HEK293 cells following transient transfection, and produced by protein A affinity chromatography followed by ion exchange chromatography as described previously.^10^ The conjugation of *p*-SCN-Bn-DOTA on YS5 was carried out according to our previously published method with slight modifications.^13^ Briefly, YS5 (5 mg) in HEPES buffer was exchanged with 0.1 M Na_2_CO_3_/NaHCO_3_ buffer (pH = 9.0) using YM30K centrifugal filter unit (Amicon ultra-0.5 mL, Regenerated cellulose, Merck Millipore Ltd.,). It was further diluted with 0.1 M Na_2_CO_3_/NaHCO_3_ buffer (pH = 9.0) to adjust the final volume to 1 mL. 20 eq. of *P*-SCN-Bn-DOTA in DMSO (4.5 mg, 20 µL) was added to the solution containing 5 mg of YS5 followed by incubation at 37 °C for 1 h. The reaction mixture was purified using a PD10 gel filtration column by eluting with 0.25 M NaOAc buffer, pH=6. The DOTA-conjugated YS5 was stored in -20 °C until the radiolabeling reactions.

For radiolabeling, 1 mCi of ^225^Ac(NO_3_)_3_ was received from Oak Ridge National Laboratory by the ^229^Th decay pathway in solid form, and it was dissolved in 0.2M HCl (100 µL). The radiolabeling was performed by incubating the 50 µCi (6 µL) of ^225^Ac(NO_3_)_3_, 2 M NH_4_OAc (50 µL, pH=5.8), L-Ascorbic acid (20 µL, 150 mg/mL), and YS5-DOTA (200 µg, 15.64 µL, 12.785 mg/mL) at 40 °C for 2 h. The radiolabeling progress was monitored by radio TLC by eluting with a mobile phase 10 mM EDTA (pH=5.5) on iTLC-SG (Gelman Science Inc., Ann Arbor, MI). The radiolabeled antibody was diluted with 300 µL of 0.9% saline, transferred to a YM30K centrifugal filtration unit (Millipore, MA, USA), and centrifuged at 10,000 rpm for 10 min. Then, 200 µL of 0.9% saline was added to the centrifugal filtration unit, followed by centrifugation at 10,000 rpm for 5 min. The [^225^Ac]DOTA-YS5 (20 µCi, 40%) conjugate was isolated, and the purity was assessed by radio-TLC [iTLC-SG 10 mM EDTA (pH=5)] immediately after purification.

To confirm the stability and aggregation of the radiolabeled antibody, Size-Exclusion Chromatography (SEC) was performed. For this, a Merck Hitachi LaChrom Elite system comprised of an L-2130 Pump, L-2200 autosampler, L-2450 DAD detector, L-2400 UV detector, and Carrol and Ramsey Associates model 105S radioactivity detector was utilized. 1X PBS buffer was used as a mobile phase using a column BioSep 5 μm SEC-s3000 290 Å (Phenomenex, Inc. 411 Madrid Ave. Torrance, CA 90501 USA) with a flow rate of 1 mL per minute. 1 ml fractions were collected and then counted on Hidex automatic gamma counter when secular equilibrium was reached at 24 h. The chromatogram was plotted as CPM Vs time. The stability of the purified radioimmunoconjugate [^225^Ac]DOTA-YS5 (0.05 mL, 7.43 µCi) was verified by incubating with human serum (0.45 mL; Sigma Aldrich) and 0.9% Saline (0.45 mL, Medline) at 37 °C at various time points (0 h, 1 day, 2 days, 3 days, 4 days, 5 days, 7 days, and 14 days). At each time point, an aliquot of 8 μl was spotted in duplicate on iTLC-SG and eluted with 10 mM EDTA (pH=5.5). The decomplexation was monitored by radio TLC and allowed for 24 h to reach secular equilibrium before counting on a BioScan Ar 2000 radio-TLC Imaging scanner.

### Target binding fraction assay using magnetic beads

The target binding fraction assay was performed according to prior protocols with modifications as described below.^22^ The vials were divided into three groups named group A (testing), group B (blocking), and group C as a control group. HisPur™ Ni-NTA magnetic beads (Catalog No.88831, Thermo Fisher Scientific) were added (40 µL) to the vials in each tube A, B, and C. Following this, 380 µL phosphate-buffered saline containing 0.05% Tween-20 (PBS-T) was added over the beads. The samples were vortexed for 15-30 seconds, and beads were trapped using DynaMag-2 magnet (Purchased from Thermo Fisher Scientific., Catalog number. 12321D) to remove the supernatant. The 5 µg of CD46 (20 µL, 250 µg/mL) and PBS-T (390 μL) were added to group A and group B, while Group C was kept as no-CD46 control. These vials were then incubated for 15 min at room temperature, and the supernatant was removed. Finally, 10 ng of the [^225^Ac]DOTA-YS5 in 1% milk PBS (10 µL) was added to groups A, B, and C. A large excess of cold YS5 (10 µg) was added for the blocking group B before adding 10 ng of [^225^Ac]DOTA-YS5. Each group was diluted with 1% milk PBS to a total 400 µL/vial volume, and vials were incubated at room temperature for 30 min. Thereafter, beads were isolated using a magnet, and the supernatant was collected in separate tubes. These beads were washed twice with 400 μL of PBS-T. The bead’s activity, supernatant, and standard (10 ng of [^225^Ac]DOTA-YS5) were measured on a Hidex gamma counter immediately (energy window 25-2000 keV), without waiting for secular equilibrium. The binding fraction percentage was determined by calculating the bead’s activity/standard.

### Cell culture

Prostate cancer (22Rv1 and DU145) and HEK293a cells were obtained from American Type Culture Collection and were cultured in a CO_2_ incubator with RPMI medium supplemented with 10% fetal bovine serum and antibiotic solution (1% penicillin and streptomycin).

### Cell Binding assay

To confirm the binding of [^225^Ac]DOTA-YS5 after radiolabeling, a binding assay was carried out, where [^225^Ac]DOTA-YS5 (7 nM) was incubated with 22Rv1 cells (3 million cells per tube, triplicate) for 1 h. The blocking samples were kept where 2-fold excess of cold YS5 was added. To avoid the non-specific binding of [^225^Ac]DOTA-YS5, 1% milk protein was added to the tubes. The cells were centrifuged (at 500 x g) and washed twice with saline to separate the unbound [^225^Ac]DOTA-YS5. The [^225^Ac]DOTA-YS5 bound to 22Rv1 cells was counted in the HIDEX gamma counter (a gamma energy window: 25 to 2000 keV), without waiting for secular equilibrium.

### Membrane-bound and internalized [^225^Ac]DOTA-YS5

The time-dependent binding and internalization of [^225^Ac]DOTA-YS5 were studied in 22Rv1 and DU145 cells. PCa cells (22Rv1 and DU145) were seeded in 24 wells plates at the density of 4×10^4^ cells per well. The cells were incubated with [^225^Ac]DOTA-YS5 (20 nM) for 30 minutes, 1 h, 2 h, 3 h and 24 h. After incubation, cells were washed with PBS and incubated with glycine buffer (50 mM glycine and 100 mM NaCl, pH 3) for 5 minutes to collect the membrane-bound fraction of [^225^Ac]DOTA-YS5. The glycine buffer-stripped membrane-bound activity was collected separately, whereas the internalized activity was organized by cell lysis with 5N NaOH solution. The membrane-bound and internalized [^225^Ac]DOTA-YS5 was counted in a gamma counter (HIDEX, energy window of 25 to 2000 keV) without attaining secular equilibrium.

### The *in-vitro* cell-killing activity of [^225^Ac]DOTA-YS5

To study the cell viability after treatment of [^225^Ac]DOTA-YS5 and non-targeted [^225^Ac]DOTA-IgG, 22Rv1 cells were seeded in 96 well plate (1K cells per well). 22Rv1 cells were treated with increasing concentrations of [^225^Ac]DOTA-YS5 or [^225^Ac]DOTA-IgG (0.0 pCi to 100 nCi/mL) for 96 h. Following the treatment, 10 µL of MTT dye (5mg/mL in PBS) was added to each well, and plates were incubated for 2 h. Thereafter, the formazan crystals formed from MTT dye were dissolved in DMSO (150 µL per well). The cell viability was measured by taking absorbance of plates at 570 nm. The %cell viability data were fitted into a sigmoidal dose-response curve using Graphpad Prism program.

### Animal studies

All the animal experiments were carried out in compliance with institutional Animal Care and Use Committee (IACUC) established guidelines at Laboratory Animal Resource Center (LARC), University of California, San Francisco. For 22Rv1 and DU145 xenografts and toxicity studies, male nude mice (4- to 6-week-old, homozygous, Nu/J, strain#: 002019) from Jackson Laboratories were used. The animals were housed in a facility with 12 h light or day cycle and free food and water access. To prepare the xenografts, 2.5 million cells in phosphate-buffered saline (PBS) were mixed with Matrigel (Corning: 356237) in a 1:1 volume ratio and were injected subcutaneously. The patient-derived xenografts (LTL-545 and LTL-484) were obtained from Living Tumor Laboratory. For LTL-545 and LTL-484, 8 to 10 week-old male NOD SCID gamma (NSG, NOD.Cg-*Prkdc*^*scid*^ *Il2rg*^*tm1Wjl*^/SzJ, Strain#:005557) mice were implanted subcutaneously with patient-derived tissue as we previously described.^13^ To euthanize the animals used in the experiments, a high dose of inhaled anesthetics (isoflurane) was given followed by cervical dislocation.

### Toxicity study in nude mice

Toxicity of the [^225^Ac]DOTA-YS5 was studied in healthy nude mice (4- to 5-week-old, homozygous) from Jackson Laboratories. For acute toxicity study, mice were administered 0.06 µCi, 0.125 µCi, 0.25 µCi, and 0.5 µCi of [^225^Ac]DOTA-YS5, or vehicle control via tail vein. For chronic toxicity, mice were injected with 0.25 µCi, and 0.5 µCi of [^225^Ac]DOTA-YS5, or vehicle control. These mice were monitored for body weight measurements (twice a week) for 15 days and 117 days for acute and chronic toxicity studies, respectively. At the end of the experiment, a complete blood count and laboratory organ function tests were carried out to study acute and chronic toxicity of the [^225^Ac]DOTA-YS5. For this, a cardiac puncture was carried out to collect blood in EDTA coated tubes to study blood cell counts. To collect the serum samples, blood containing vials were allowed to sit at 4 °C for 30 min. The clotted blood samples were centrifuged (15,000 rpm, 10 min, at 4 °C) to separate the serum. Blood and serum samples were sent to the Comparative Pathology Laboratory, UC Davis School of Veterinary Medicine for blood cell counts and organ function tests.

### Histology of the tissue sections

Following euthanasia, the organs (liver, kidney, lungs, spleen, heart, and brain) were removed from all animals and fixed in 10% neutral buffered formalin. Routine histologic analysis was performed to study microscopic features of the tissues. For H&E, tissues were formalin-fixed, processed with ethanol gradient (from 30% to 70%), and paraffin-embedded. The tissue sections of 4 µm thickness were processed for H&E staining. Periodic acid-Schiff (PAS) and Trichrome staining were performed on a subset of organs as needed. [^225^Ac]DOTA-YS5 related toxicity was assessed by comparing histologic changes in treated mice relative to saline control mice.

### Immunofluorescence for DNA damage analysis

The 22Rv1 xenograft-bearing mice were injected with saline or 0.5 µCi dose of the [^225^Ac]DOTA-YS5. Tumor tissues were collected after 7 days p.i. and 14 days p.i., followed by fixation (10% formalin) and sectioning. The sections were stained with H&E, DAPI (nuclear stain) and anti-phospho-γ-H2AX (CST #9718) antibodies. The phospho-γ-H2AX foci as well as the nuclear areas were analyzed using ImageJ program.

### [^225^Ac]DOTA-YS5 Biodistribution

To study the biodistribution of [^225^Ac]DOTA-YS5, 22Rv1 xenograft-bearing mice were used. These mice with tumor volumes ranging from 200-300 mm^3^ received a 0.5 µCi dose of [^225^Ac]DOTA-YS5 via tail vein injections. Following this, the blood, tumor, and selected organs were collected after 24 h (Day 1), 48 h (Day 2), 96 h (Day 4), 168 h (Day 7), 264 h (Day 11), 336 h (Day 14), and 408 h (Day 17) to study activity present in the tissue. Following this, the presence of [^225^Ac]DOTA-YS5 in each organ was counted (gamma energy window: 25 to 2000 keV) with the gamma counter (HIDEX). The standards of injected dose were measured in a similar fashion to calculate the %ID/g for each organ without reaching secular equilibrium.

### iQID camera alpha-particle imaging

To study the distribution of [^225^Ac]DOTA-YS5 within the tissue, iQID (ionizing radiation quantum imaging detector) camera imaging was performed. For this, tumor, liver, kidneys, spleen, heart, and lungs tissues from the mice injected with a 0.5 µCi dose of [^225^Ac]DOTA-YS5 were collected in an Optimum Cutting Temperature (OCT) medium. The tissues in OCT were sliced (20 µm thick) and mounted with a scintillator on the iQID camera (QScint Imaging Solutions, LLC) and acquisition of alpha particles was done for 24h at 1.5V.

### Dosimetry

Dosimetry calculations were performed by using the time-integrated activity coefficients with curve-fitting performed in the EXM module of OLINDA/EXM Version 1.1, and the digital mouse phantom in OLINDA Version 2.0.^43^ For this, the data from the biodistribution study was organized to produce time and % of injected activity for each organ and tumor. Following this, these numbers were used as inputs for deriving time-integrated activity coefficients, and calculating equivalent dose (in Sv) that is a product of absorbed dose (in Gy) and radiation weighting factors.

### Treatment studies with [^225^Ac]DOTA-YS5

For all treatment studies, the body weights and tumor measurements for each mouse were measured until the mice reached a humane endpoint including body condition score below 2, weight loss by 20%, or tumor volume of 2000 mm^3^. The tumor measurements were carried out twice a week, and tumor volumes were calculated by formula V=[length*(width)^2^]/2.

Animals were randomized in groups (n=7), and [^225^Ac]DOTA-YS5 injections were performed when the tumor reached 100 mm^3^. The 22Rv1 and DU145 xenograft bearing mice were injected with saline, 0.25 µCi, or 0.5 µCi of [^225^Ac]DOTA-YS5. Additionally, a separate group received a 0.5 µCi dose of [^225^Ac]DOTA-IgG. For fractionated dose study, the treatment group was administered with three doses of 0.125 µCi of [^225^Ac]DOTA-YS5 on day 1, day 37, and day 49. The LTL-545 bearing mice were injected with saline and [^225^Ac]DOTA-YS5 (0.03, 0.06, 0.12, and 0.25 µCi) along with native IgG (0.5 mg per mouse) for Fc blocking. Similarly, the LTL-484 mice were injected intravenously with 0.06 µCi, 0.12 µCi, and 0.25 µCi doses of [^225^Ac]DOTA-YS5 when tumors reached to 100 mm^3^.

### Statistical analysis

The results reported are presented as Mean ±SD and plotted using Graphpad Prism software. To compare the blood cells and liver and kidney function tests between saline control and treated groups one way ANOVA was used with Dunnett’s multiple comparisons test. The significance of median survival from saline control and treatment groups was determined by Log-rank (Mantel-Cox) test. Students t-test was used to compare the number of phospho-γ-H2AX foci in sections from saline and [^225^Ac]DOTA-YS5 treated mice groups.

## Acknowledgments

RRF acknowledges funding from Translational Science awards from the Prostate Cancer Research Program of the Congressionally Directed Medical Research Program of the US Department of Defense (W81XWH-21-1-0792, W81XWH-20-1-0292), Pilot funding through the UCSF Precision Imaging of Cancer and Therapy Program, and pilot funding through the UCSF Cancer Imaging Resource and Preclinical Therapeutics Core through P30CA082103. ^225^Ac was supplied by the U.S. Department of Energy Isotope Program, managed by the Office of Isotope R&D and Production. MALDI-MS data were provided by the Mass Spectrometry Facility, Department of Chemistry, University of Alberta (Edmonton, Alberta, Canada). Blood cell counts and organ function tests were performed by Comparative Pathology Laboratory, UC Davis School of Veterinary Medicine. Histology staining procedures were carried out by Histology and Biomarker core, University of California San Francisco.

## Competing interests

R.R.F. reports prior grant funding from Fukushima SiC, outside the existing work.

BL is a founder, board member and equity holder of Fortis Therapeutics, Inc., which licensed intellectual properties from the University of California and is conducting clinical trials on CD46 targeting agents. BL also holds equity of Molecular Imaging and Therapeutics, Inc., which were converted to equity of Fortis Therapeutics. BL is an inventor of intellectual properties around the CD46 epitope, the CD46 targeting human antibody, and therapeutic targeting of CD46.

SB is an inventor of intellectual properties around the CD46 epitope, the CD46 targeting human antibody, and therapeutic targeting of CD46, which were licensed to Fortis Therapeutics, Inc.

YSu is an inventor of intellectual properties around the CD46 epitope, the CD46 targeting human antibody, and therapeutic targeting of CD46, which were licensed to Fortis Therapeutics, Inc.

JH holds equity of Molecular Imaging and Therapeutics, Inc., which were converted to equity of Fortis Therapeutics that licensed intellectual properties from the University of California.

YSeo holds equity of Molecular Imaging and Therapeutics, Inc., which were converted to equity of Fortis Therapeutics that licensed intellectual properties from the University of California.

## Author contributions

A.P.B. performed *in vitro* and *in vivo* studies, analyzed the data, and wrote the manuscript. S.W. optimized radiolabeling conditions and contributed to µPET scan and therapeutic efficiency studies. K.N.B. performed radiochemistry experiments, and contributed to in vivo studies, manuscript writing, and data analysis. E.C. analyzed and interpreted and wrote the results from histology experiments. S.B. performed antibody purification and characterization, and data analysis. E.A.E and J.C. performed western blotting experiments, provided the PDX tumor models, and participated in data analysis. R.P. provided guidance in performing alpha particle imaging. U.A. helped in chronic toxicity studies. N.M. provided guidance and performed data analysis, A.A. participated in biodistribution studies and data analysis. S.D. participated in vivo experiments and data analysis. D.B.V. participated in data analysis and provided guidance in radiolabeling reactions. Y.S. optimized the YS5 antibody purification process and participated in data analysis. R.T. helped in imaging µPET/CT scan experiments. J.H. provided experimental guidance and participated in data analysis. D.M.W. provided experimental guidance and participated in data analysis. L.Z. performed statistical analyses. R.A. provided guidance for experimental design and participated in data analysis. H.F.V provided guidance for experimental design and provided important revisions. B.L. conceptualized the study, provided support, guidance for experimental design, data analysis and manuscript drafting and revision. R.R.F. conceptualized the study, designed, funded, guided the project, and wrote the manuscript. All the authors have contributed and corrected the manuscript.

