## Supplementary Information for "Treatment of prostate cancer with CD46 targeted ^225^Ac alpha particle radioimmunotherapy"

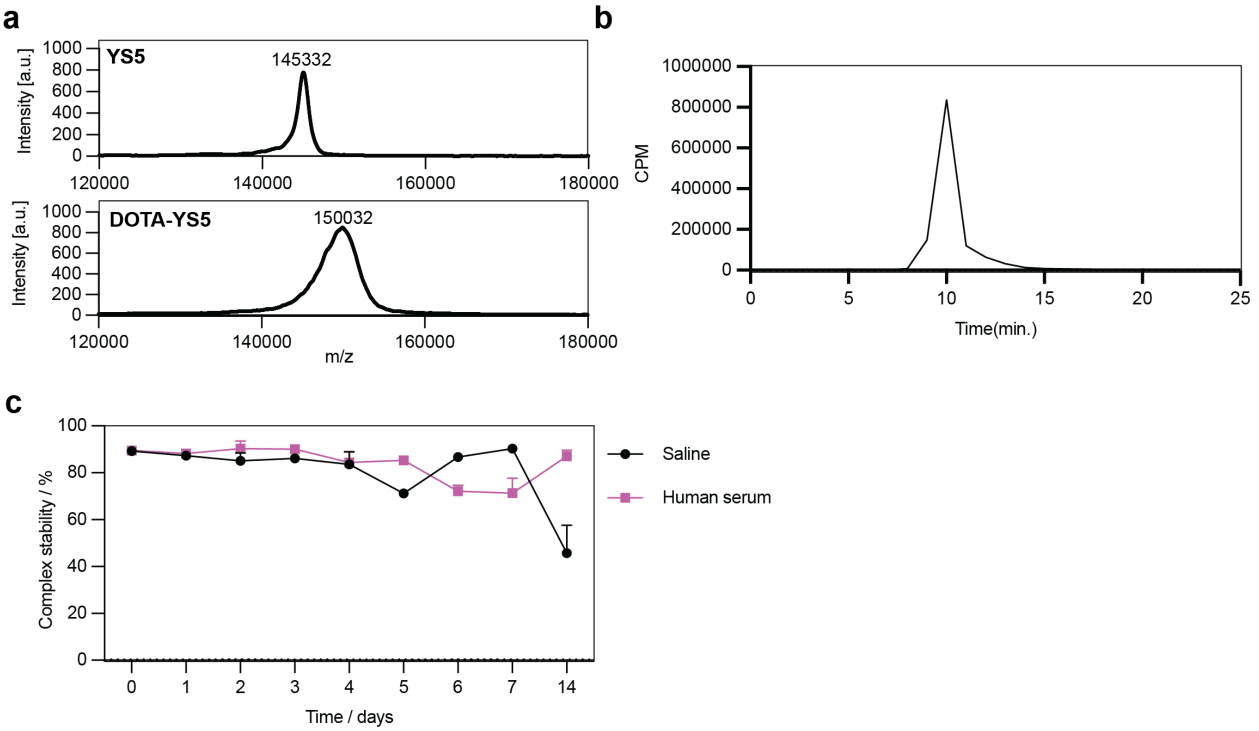


**Figure S1**: **a** MALDI-TOF Mass spectroscopy results show 8.7 equivalents of DOTA molecules on the YS5 antibody. The number of DOTA were calculated by dividing the difference of m/z between YS5 and DOTA-YS5 by the molecular weight of DOTA. **b** Size-exclusion chromatogram shows no aggregation of the [$^{225}$Ac]DOTA-YS5 after the radiolabeling steps. **c** Stability of the [$^{225}$Ac]DOTA-YS5 tested in saline and human serum for 14 days.

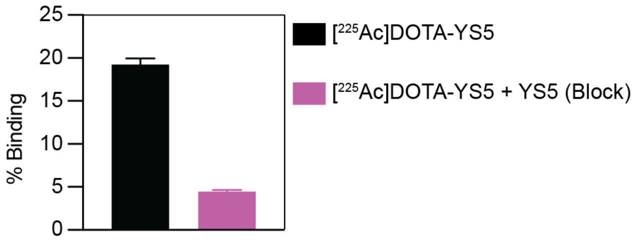


**Figure S2**: Binding of [$^{225}$Ac]DOTA-YS5 to 22Rv1 with or without cold antibody blocking. Blocking with cold YS5 results in a reduction of binding from 19.25±0.70% to 4.46±0.17%.

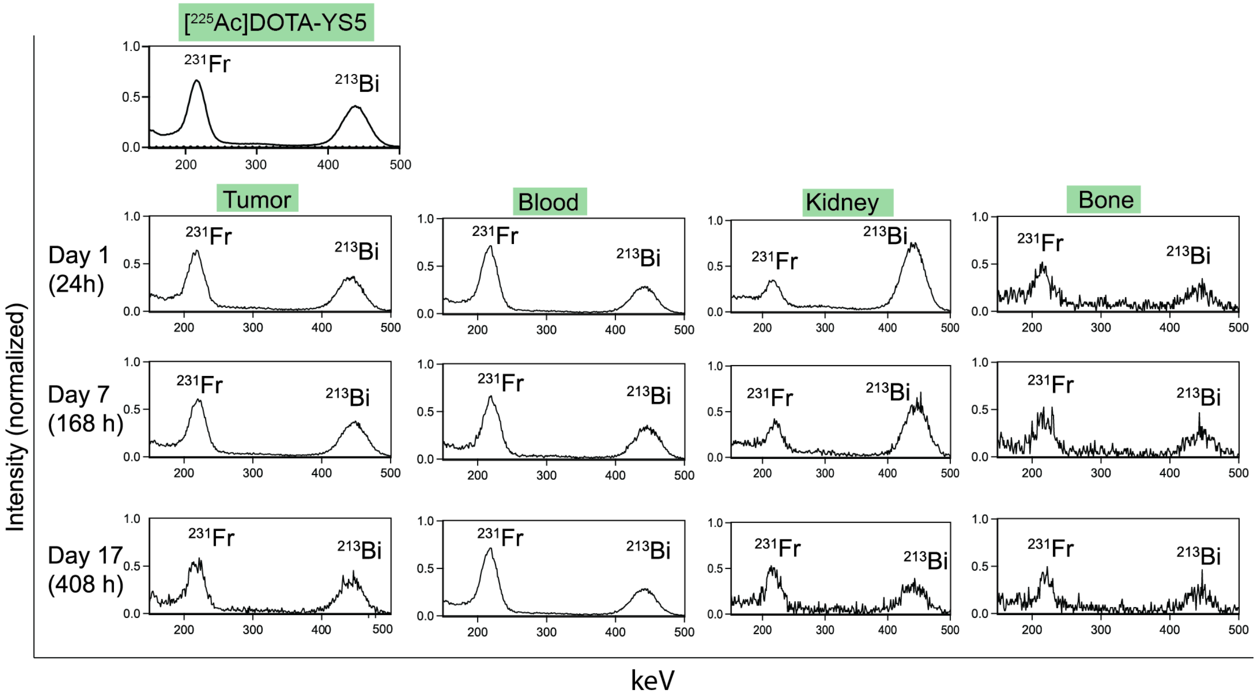


**Figure S3**: Gamma energy spectra of the tumor, blood, kidneys, and bone showing the intensities of the $^{213}$Bi and $^{221}$Fr peaks. As compared to the equilibrium gamma energy spectra of [$^{225}$Ac]DOTA-YS5, increased intensity of the $^{213}$Bi was observed in kidney samples.

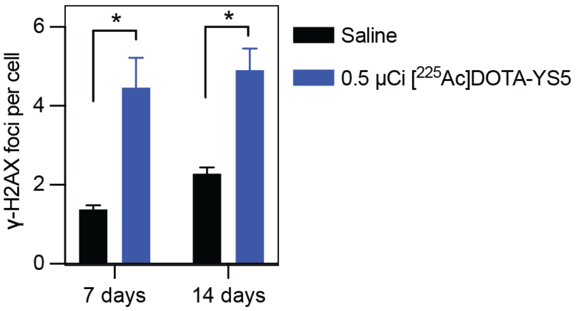


**Figure S4**: Number of H2AX-foci after the treatment of saline and [$^{225}$Ac]DOTA-YS5 for 7 days and 14 days. t-test *p* value is indicated as * $p<0.05$.

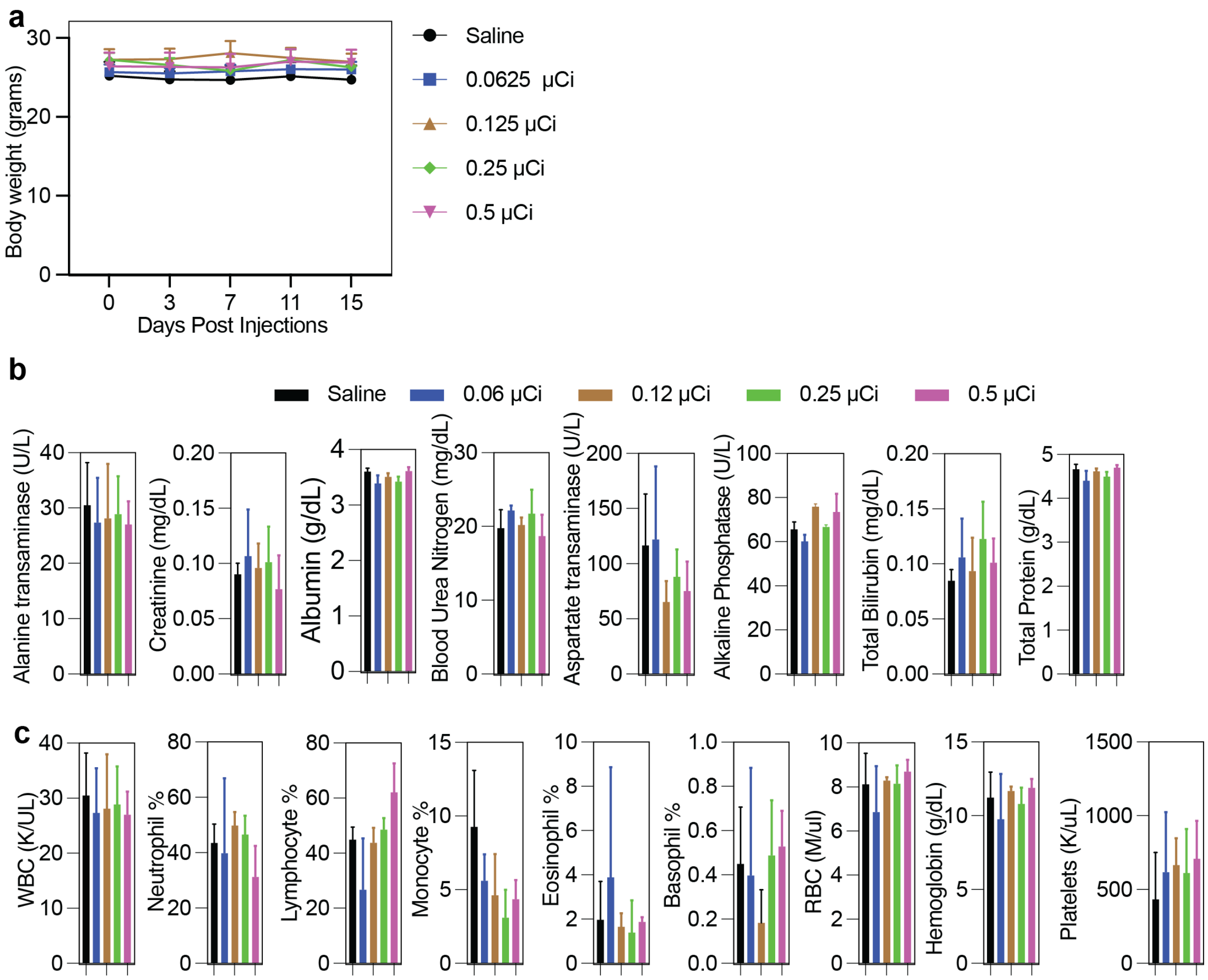


**Figure S5:** Acute toxicity study of the [$^{225}$Ac]DOTA-YS5 in nude mice. **a** The body weights of mice injected with [$^{225}$Ac]DOTA-YS5 showed no significant change in treatment groups over the 15 days study period. **b&c**: Liver and kidney function tests (**b)** and blood parameters (**c**) show no significant changes for saline vs. treated groups indicating the short-term safety of [$^{225}$Ac]DOTA-YS5.

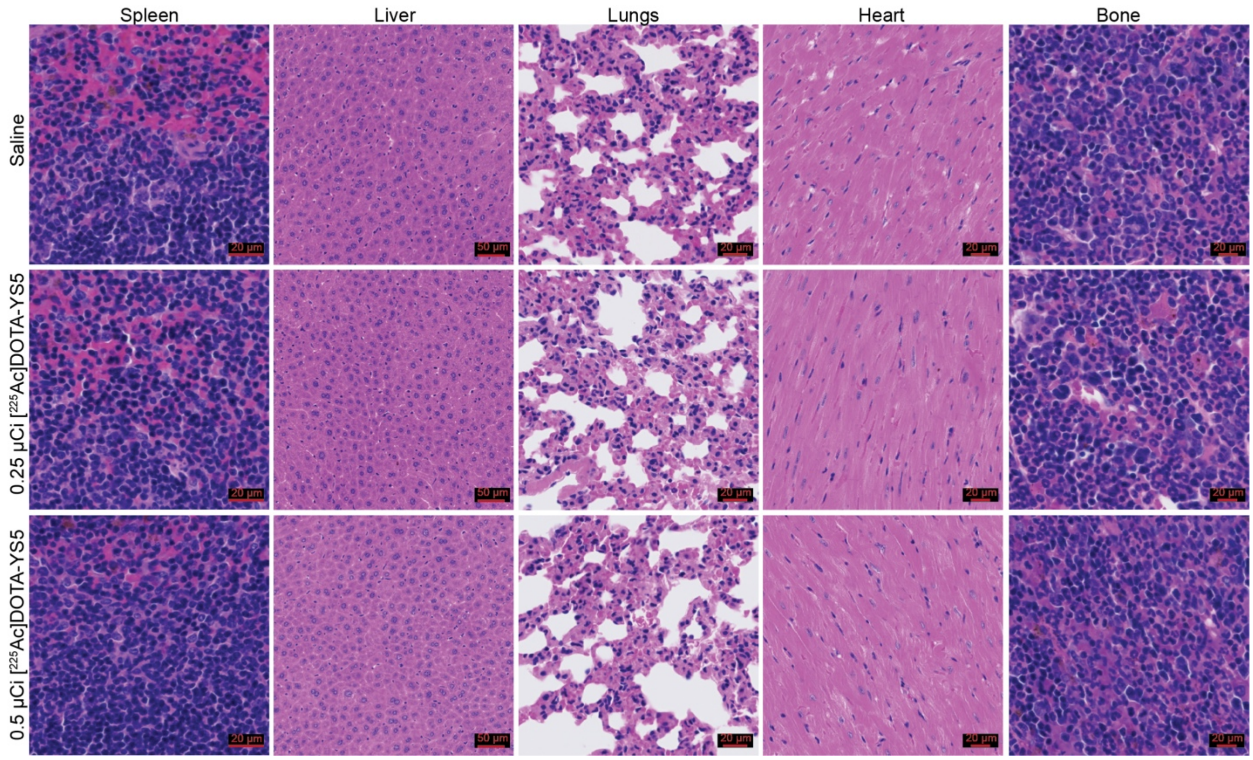


**Figure S6**: Histology evaluation of the healthy tissues for long-term (117 days) toxicity analysis. Hematoxylin and Eosin (H&E) staining of spleen, liver, lungs, heart, and bone samples show no toxicity at 0.25 µCi or 0.5 µCi doses of [$^{225}$Ac]DOTA-YS5. Scale bar: 20 µm.

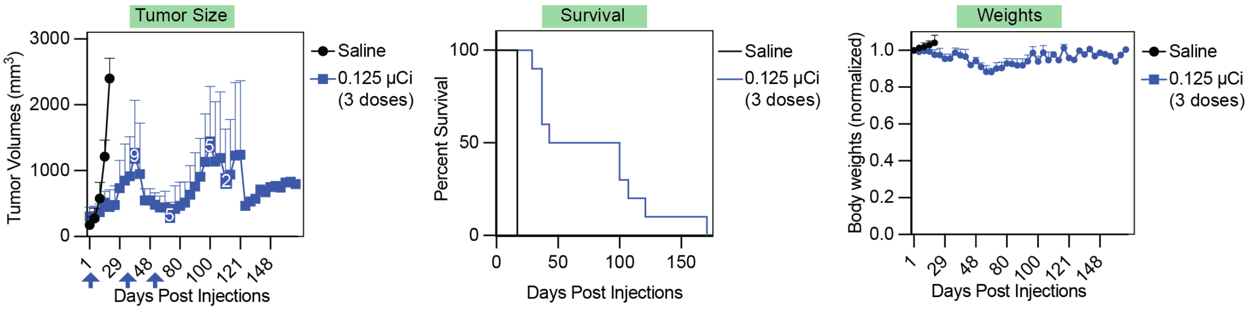


**Figure S7**: Tumor volumes, overall survival, and body weights for the saline and fractionated dose (0.125 μCi x 3) injections in 22Rv1 xenografts. Results show delayed tumor growth and improved survival without the significant toxicity from fractionated dose treatment.

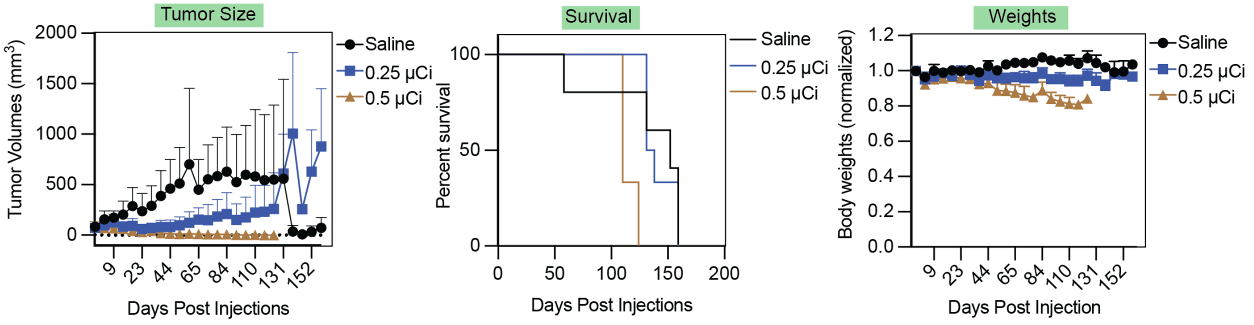


**Figure S8**: Antitumor activity of [$^{225}$Ac]DOTA-YS5 in subcutaneous DU145 tumors in nude mice. Tumor volume, survival analysis, and weight measurements after the injections of [$^{225}$Ac]DOTA-YS5 in DU145 xenografts.

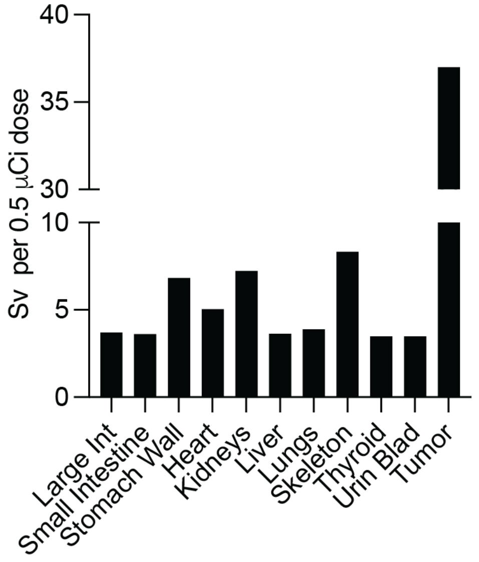


**Figure S9**: Estimated equivalent doses (in Sv) in organs and tumor indicating the highest dose (37 Sv) was delivered to tumor tissue.

**Table S1**: Biodistribution of $^{68}$Ga-PSMA11 Patient-derived xenograft models of prostate cancer (LTL-545, LTL-484, LTL-331, and LTL-331R). Data is shown as mean %ID/gm ± SD. (n = 4)

| | LTL-545 | LTL-484 | LTL-331 | LTL-331R |
|---|---|---|---|---|
| Blood | 0.13±0.06 | 0.08±0.01 | 0.17±0.02 | 0.18±0.06 |
| Tumor | 3.32±0.41 | 4.41±1.11 | 2.3±0.83 | 0.44±0.15 |
| Muscle | 0.46±0.12 | 0.21±0.06 | 0.42±0.15 | 0.33±0.11 |
| Spleen | 23.1±5.06 | 7.67±0.73 | 19.36±5.95 | 18.64±4.19 |
| Pancreas | 0.89±0.13 | 0.46±0.07 | 0.85±0.17 | 0.78±0.22 |
| Kidney | 137.25±13.01 | 113.55±19.91 | 127.77±17.39 | 138.78±16.78 |
| Liver | 0.37±0.12 | 0.32±0.1 | 0.32±0.09 | 0.47±0.15 |
| Lung | 2.64±0.34 | 1.42±0.36 | 2.23±0.6 | 2.95±0.54 |
| Bone | 0.28±0.1 | 0.09±0.03 | 0.36±0.28 | 0.17±0.06 |

**Table S2**: [$^{225}$Ac]DOTA-YS5 biodistribution data in 22Rv1 xenograft mice for day 1 to day 17 (mean %ID/gm ± SD). (n = 4)

| Organ | Day 1 | Day 2 | Day 4 | Day 7 | Day 11 | Day 14 | Day 17 |
|---|---|---|---|---|---|---|---|
| **Blood** | 11.64±1.37 | 9.46±1.63 | 9.8±1.8 | 3.94±0.95 | 1.88±0.77 | 1.87±0.45 | 1.71±0.47 |
| **Tumor** | 11.17±4.76 | 17.49±10.85 | 28.58±10.93 | 29.35±7.76 | 18.1±8.12 | 21.11±12.3 | 31.76±5.9 |
| **Muscle** | 0.4±0.01 | 1.08±0.08 | 0.58±0.33 | 0.39±0.1 | 0.12±0.07 | 0.22±0.14 | 0.14±0.09 |
| **Spleen** | 2.7±0.64 | 7.4±1.37 | 4.01±0.78 | 3.05±0.57 | 1.7±1.03 | 1.32±0.44 | 2.19±0.97 |
| **Pancreas** | 0.68±0.12 | 0.94±0.24 | 0.58±0.13 | 0.41±0.03 | 0.27±0.05 | 0.27±0.09 | 0.27±0.18 |
| **Kidneys** | 12.78±1.85 | 4.48±0.83 | 3.6±0.72 | 3.29±0.79 | 2.78±0.94 | 1.65±0.43 | 1.49±0.31 |
| **Stomach** | 0.96±0.44 | 0.59±0.1 | 0.55±0.16 | 0.5±0.07 | 0.22±0.1 | 0.2±0.1 | 0.27±0.03 |
| **Large Intestine** | 1.12±0.48 | 1.29±0.45 | 1.17±0.28 | 0.48±0.2 | 0.47±0.17 | 0.43±0.12 | 0.46±0.22 |
| **Small Intestine** | 1.73±0.38 | 1.6±0.94 | 0.76±0.11 | 0.81±0.16 | 0.33±0.14 | 0.37±0.15 | 0.25±0.17 |
| **Liver** | 6.61±1.1 | 6.54±2.74 | 5.03±0.67 | 10.32±2.96 | 6.08±2.57 | 5.27±2.08 | 5.14±1.37 |
| **Lungs** | 3.21±0.66 | 5.63±0.51 | 2.76±0.76 | 2.23±0.9 | 1±0.78 | 1.25±0.33 | 1.31±0.42 |
| **Heart** | 4.94±0.7 | 3.65±0.94 | 3.7±0.91 | 1.82±0.45 | 1.15±0.45 | 0.84±0.17 | 0.69±0.1 |
| **Brain** | 0.14±0.07 | 0.22±0.06 | 0.22±0.12 | 0.19±0.06 | 0.1±0.05 | 0.13±0.07 | 0.07±0.04 |
| **Bone** | 3.1±0.46 | 5.78±1.53 | 5.7±1.71 | 5.65±2.01 | 4.82±1.93 | 10.17±5.74 | 6.87±0.53 |

**Table S3**: Tumor to blood, tumor to muscle and tumor to kidney ratios of [$^{225}$Ac]DOTA-YS5 at Day 1 to day 17. (Mean ± SD, n = 4)

| | Tumor to blood | Tumor to muscle | Tumor to kidneys |
|---|---|---|---|
| **Day 1** | 0.96±0.37 | 27.71±11.3 | 0.89±0.45 |
| **Day 2** | 1.85±1.02 | 16.67±10.81 | 3.92±2.18 |
| **Day 4** | 3.06±1.47 | 64.92±42.17 | 8.51±4.27 |
| **Day 7** | 7.98±3.33 | 81.96±41.61 | 9.38±3.67 |
| **Day 11** | 9.45±2.62 | 139.76±13.27 | 6.45±2.67 |
| **Day 14** | 11.21±4.74 | 176.95±204.69 | 13.33±6.99 |
| **Day 17** | 19.37±4.55 | 347.33±249.41 | 21.96±5.27 |

**Table S4**: Liver and kidney function test results for the acute toxicity study group. (Mean ± SD, n = 5)

| | Saline | [$^{225}$Ac]DOTA-YS5 | | | |
|---|---|---|---|---|---|
| | | 0.06 µCi | 0.125 µCi | 0.25 µCi | 0.5 µCi |
| **Albumin g/dL** | 3.6±0.05 | 3.39±0.13 | 3.50±0.06 | 3.425±0.09 | 3.61±0.07 |
| **Total Protein g/dL** | 4.66±0.10 | 4.39±0.20 | 4.6±0.05 | 4.49±0.11 | 4.7±0.06 |
| **Blood Urea Nitrogen mg/dL** | 19.75±0.10 | 22.17±0.57 | 20.10±1.13 | 21.72±3.26 | 18.7±2.88 |
| **Creatinine mg/dL** | 0.09±0.10 | 0.10±0.04 | 0.09±0.02 | 0.10±0.03 | 0.07±0.03 |
| **Total Bilirubin mg/dL** | 0.08±0.01 | 0.10±0.03 | 0.09±0.03 | 0.12±0.03 | 0.10±0.02 |
| **Alanine transaminase U/L** | 30.5±6.69 | 27.35±7.02 | 28.1±8.57 | 28.87±6.88 | 27±4.20 |
| **Aspartate transaminase U/L** | 116.37±40.48 | 122.05±57.24 | 65.07±16.37 | 88.05±24.79 | 75.1±26.73 |
| **Alkaline Phosphatase U/L** | 65.6±2.89 | 60.175±2.56 | 75.85±1.19 | 66.725±0.96 | 73.5±8.21 |

**Table S5**: Blood parameters for the acute toxicity study groups (MCV: mean corpuscular volume, MCH: mean corpuscular hemoglobin, , RDW: red cell distribution width, MPV: mean platelet volume). (Mean ± SD, n = 5)

| | Saline | [$^{225}$Ac]DOTA-YS5 | | | |
|---|---|---|---|---|---|
| | | **0.06 μCi** | **0.125 μCi** | **0.25 μCi** | **0.5 μCi** |
| **Neutrophil %** | 43.43 ± 6.90 | 39.77 ± 27.12 | 49.80 ± 4.87 | 46.48±6.89 | 31.19 ± 11.21 |
| **Lymphocyte %** | 44.87 ± 4.62 | 26.73 18.57 | 43.74 ± 5.44 | 48.55±4.11 | 62.05 ± 10.56 |
| **Monocyte %** | 9.26 3.83 | 4.20 ± 3.16 | 4.62 ± 2.79 | 3.09±1.89 | 4.35 ± 1.30 |
| **Eosinophil %** | 1.97 ± 1.73 | 3.88 ± 4.97 | 1.64 ± 0.62 | 1.38±1.47 | 1.87 ± 0.21 |
| **Basophil %** | 0.45 ± 0.25 | 0.39 ± 0.048 | 0.18 ± 0.14 | 0.48±0.25 | 0.52 ± 0.16 |
| **WBC (K/ul)** | 0.77 ± 0.22 | 1.45 ± 1.08 | 1.25 ± 0.34 | 1.37 ±0.33 | 2.96 ± 3.80 |
| **Absolute Neutrophil cells (K/ul)** | 0.34 ± 0.13 | 0.72 ± 0.54 | 0.62 ±0.22 | 0.63±0.13 | 0.692 0.62 |
| **Absolute Lymphocyte cells (K/ul)** | 0.34 ± 0.09 | 0.49 ± 0.39 | 0.54 ± 0.15 | 0.67±0.21 | 2.05 ± 2.88 |
| **Absolute Monocyte cells (K/ul)** | 0.07 ± 0.02 | 0.08 ± 0.08 | 0.05 ± 0.02 | 0.04±0.03 | 0.15 ± 0.22 |
| **Absolute Eosinophil cells (K/ul)** | 0.01 ± 0.00 | 0.09 ± 0.14 | 0.01 7 ±0.005 | 0.02±0.02 | 0.05 ± 0.06 |
| **Absolute Basophil cells (K/ul)** | 0.002 ± 0.005 | 0.007 ± 0.015 | 0 | 0.005±0.01 | 0.01± 0.01 |
| **Platelets (K/uL)** | 433.25 ± 316.79 | 616.25 ± 407.20 | 665.25 ± 181.89 | 612.25±297.23 | 707.25 ± 257.89 |
| **RBC (M/ul)** | 8.11 ± 1.40 | 6.85 ± 2.09 | 8.28 ± 0.15 | 10.8±1.10 | 8.69 ± 0.53 |
| **Hemoglobin (g/dL)** | 11.22 ± 1.73 | 9.75 ± 3.07 | 11.67 0.29 | 36.42±3.79 | 11.87 ± 0.61 |
| **Hematocrit %** | 37.95 ± 6.35 | 31.27 10.07 | 37.6 ± 0.85 | 612.25±297.23 | 39.22 ± 2.46 |
| **MCV (fL)** | 46.77 ± 0.70 | 45.37 ± 1.17 | 45.37 ± 0.72 | 45.10 ± 0.14 | 44.7±0.32 |
| **MCH (pg)** | 13.85 ± 0.52 | 14.2 ± 0.50 | 14.1 ± 0.18 | 13.67 ± 0.12 | 13.3±0.45 |
| **RDW %** | 19.30 ± 0.40 | 18.72 ± 1.25 | 19.12 ± 0.12 | 19.20 ± 0.25 | 18.82±0.30 |
| **MPV (fL)** | 5.45 ± 0.52 | 5.55 ± 0.17 | 5 ± 0.08 | 5.10 ± 0.35 | 5.05±0.13 |

**Table S6**: Liver and kidney function tests results for chronic toxicity group. (Mean ± SD, n = 5)

| | **Saline** | **[$^{225}$Ac]DOTA-YS5** | |
|---|---|---|---|
| | | **0.25 µCi** | **0.5 µCi** |
| **Alanine transaminase U/L** | 20.4± 2.3 | 18± 4.4 | 28.5± 12.7 |
| **Aspartate transaminase U/L** | 70.2± 22 | 70.3± 18.6 | 164± 135.9 |
| **Albumin g/dL** | 3.0± 0.2 | 3.0 ± 0.5 | 3.8± 0.1 |
| **Alkaline Phosphatase U/L** | 50.7± 7.8 | 82.8± 6.8 | 178.8± 34.7 |
| **Blood Urea Nitrogen mg/dL** | 24.7± 1.9 | 28.7± 1.7 | 58.9± 13.5 |
| **Creatinine mg/dL** | 0.1± 0 | 0.1± 0 | 0.3± 0.1 |
| **Total Bilirubin mg/dL** | 0.1± 0 | 0.1± 0 | 0.1± 0 |
| **Total Protein g/dL** | 4.5± 0.2 | 4.4± 0.5 | 5.2± 0 |

**Table S7**: Blood parameters for the chronic toxicity study group (MCV: mean corpuscular volume, MCH: mean corpuscular hemoglobin, MCHC: mean corpuscular hemoglobin concentration, RDW: red cell distribution width, MPV: mean platelet volume). (Mean ± SD, n = 5)

| | Saline | [$^{225}$Ac]DOTA-YS5 | |
|---|---|---|---|
| | | 0.25 µCi | 0.5 µCi |
| WBC (K/ul) | 2.3± 1 | 1.6± 0.9 | 2.1± 1.5 |
| Absolute Neutrophil cells (K/ul) | 0.9± 0.6 | 0.6± 0.4 | 0.7± 0.7 |
| Absolute Lymphocyte cells (K/ul) | 1.2± 0.6 | 0.9± 0.5 | 0.9± 0.5 |
| Absolute Monocyte cells (K/ul) | 0.2± 0 | 0.1± 0.1 | 0.3± 0.2 |
| Absolute Eosinophil cells (K/ul) | 0± 0 | 0± 0 | 0.1± 0.2 |
| Absolute Basophil cells (K/ul) | 0± 0 | 0± 0 | 0± 0.1 |
| RBC (M/ul) | 6.1± 1.9 | 5.7± 1.8 | 6.1± 1.1 |
| Hemoglobin (g/dL) | 8.1± 2.3 | 7.6± 2.5 | 7.4± 1.4 |
| Hematocrit % | 27.1± 8.2 | 25.5± 8.2 | 26.5± 5.1 |
| Neutrophil % | 36.6± 10.2 | 31.2± 11 | 25.1± 12 |
| Lymphocyte % | 51.4± 11.1 | 57.7± 7.8 | 52.7± 15.4 |
| Monocyte % | 10.2± 6 | 9.6± 3.3 | 16.9± 6.4 |
| Eosinophil % | 1.3± 0.2 | 1.1± 0.6 | 4.2± 4 |
| Basophil % | 0.5± 0.1 | 0.4± 0.1 | 1.1± 1.3 |
| MCV (fL) | 44.8± 0.6 | 44.8± 0.6 | 43.5± 0.7 |
| MCH (pg) | 13.4± 0.3 | 13.4± 0.3 | 12.2± 0.2 |
| MCHC (g/dL) | 29.8± 0.7 | 29.9± 0.4 | 27.9± 0.2 |
| RDW % | 18.8± 0.4 | 18.6± 0.4 | 19± 0.4 |
| Platelets (K/uL) | 820± 601.7 | 571.7± 403.5 | 641.7± 427 |
| MPV (fL) | 5.1± 0.6 | 5.1± 0.3 | 5.6± 0.8 |
